# Anthroponumbers.org: A Quantitative Database Of Human Impacts on Planet Earth

**DOI:** 10.1101/2022.03.04.483053

**Authors:** Griffin Chure, Rachel A. Banks, Avi I. Flamholz, Nicholas S. Sarai, Mason Kamb, Ignacio Lopez-Gomez, Yinon Bar-On, Ron Milo, Rob Phillips

## Abstract

The Human Impacts Database (www.anthroponumbers.org) is a curated searchable resource housing quantitative data relating to the diverse anthropogenic impacts on our planet, with topics ranging from sea level rise, to livestock populations, greenhouse gas emissions, fertilizer use, and beyond. Each entry in the database relates a quantitative value (or a time-series of values) along with clear referencing of the primary source, the method of measurement or estimation, an assessment of uncertainty, links to the underlying data, as well as a permanent identifier called an Human Impacts ID (“HuID”). While there are other databases that house some of these values, they are typically focused on a single topic area like energy usage or greenhouse gas emissions. The Human Impacts Database provides centralized access to quantitative information about the myriad ways in which humans impact the Earth, giving links to more specialized databases for interested readers. Here, we outline the structure of the database and describe our curation procedures. Finally, we use this database to generate a graphical summary of the current state of human impacts on the Earth, illustrating both their numerical values and their dense interconnections.

**The Bigger Picture:** Over the last 10,000 years, human activities have transformed the Earth through farming, forestry, mining and industry. The complex results of these activities are now observed and quantified as “human impacts” on Earth’s atmosphere, oceans, biosphere and geochemistry. While myriad studies have explored facets of human impacts on the planet, they are necessarily technical and often tightly-focused. Thus, finding reliable quantitative information requires a significant investment of time to assess each quantity, its methods of determination, and associated uncertainty. We present the Human Impacts Database (www.anthroponumbers.org), which houses a diverse array of such quantities. We review a subset of these values and how they help build intuition for understanding the Earth-human system. While collation alone does not tell us how to best ameliorate human impacts, we contend that any future plans should be made in light of a quantitative understanding of the interconnected ways in which humans impact the planet.

## Introduction

One of the most important scientific developments of the last two centuries is the realization that the evolution of the Earth is deeply intertwined with the evolution of life. Perhaps the most famous example of this intimate relationship is the large-scale oxygenation of Earth’s atmosphere following the emergence of photosynthesis^1^. This dramatic change in the composition of the atmosphere is believed to have caused a massive extinction, as the organisms living at the time were not adapted to an oxygenated atmosphere^2–4^. Over the past 10,000 years, humans have likewise transformed the planet, directly affecting the rise and fall of ecosystems^5–13^, the pH and surface temperature of the oceans^14,15^, the composition of terrestrial biological and human-made mass^16,17^, the planetary albedo and ice cover^18–27^, and the chemistry of the atmosphere^28–33^, to name just a few examples. The breadth of human impacts on the planet is so diverse that it touches on nearly every facet of the Earth system and every scientific discipline.

Technological advances in remote sensing, precision measurement, and computational power have made it possible to measure these anthropogenic impacts with unprecedented depth and resolution. However, as scientists with different training use distinct methods for measurement and analysis, report data in different units and formats, and use nomenclature differently, these studies can be very challenging to understand and relate to one another. Even seemingly simple questions such as “how much water do humans use?” can be difficult to answer when search engines are not optimized for finding numeric data, and a search of the scientific literature yields an array of complicated analyses with different units, varying definitions about what constitutes water use, and distinct approaches to quantifying flows. This problem persists beyond the primary scientific literature as governmental, intergovernmental, and industry datasets can be similarly cryptic and laborious to interpret.

Writing from California, as several of the authors are, where we now have a “wildfire season” and a multi-decadal drought^34,35^, we wanted to develop a deeper understanding of the ways in which human activities might have produced such dramatic and consequential changes in our local and global environment. In pursuit of basic understanding, we asked many questions like “how much water and land do humans use?” and “how much methane is emitted annually?” In our search for answers, even when the question is well defined, as is the case for methane emissions, we often encountered the same challenges: disparate technical studies written for expert audiences must be understood, evaluated and synthesized just to answer simple questions. It seemed to us that a referenced compendium of “things we already know” akin to the CRC Handbook of Chemistry and Physics would be very useful for us and others.

In building the Human Impacts Database, we took inspiration from our previous experience building and using the BioNumbers Database^36^ (bionumbers.hms.harvard.edu), a compendium of quantitative values relating to cell and organismal biology. Over the past decade, the BioNumbers Database has become a widely-accessed resource that serves not only as an index of biological numbers, but also as a means of finding relevant primary literature, learning about methods of measurement, and teaching basic concepts in cell biology^37^. We believe that a reference for quantitative data about the extent of human impacts on our planet would be similarly transformative for researchers, students and interested non-scientists. While reading an entry in the Human Impacts Database is not a replacement for reading the primary literature, exploring the various quantities the repository houses can reveal a great deal about our planet, the human civilizations living on it, and their collective impacts on the Earth. We do not know which approaches to remediating these impacts are most efficient, expedient or cost-effective, but we are convinced that proposals should be evaluated in the light of a comprehensive and quantitative understanding of the Earth-human system.

## Results

### Finding and compiling numbers from scientific literature, governmental and non-governmental reports, and industrial datasets

We have established the Human Impacts Database (http://anthroponumbers.org) as a repository for the rapid discovery of quantities describing the Earth-human system. We here provide a more complete description of the database structure, the values it holds, and the stories it tells us about how humans impact the Earth. As of this writing, the database holds > 300 unique manually-curated entries covering a breadth of data sources, including primary scientific literature, governmental and non-governmental reports, and industrial communiques. Before it is added to the database and made public, each entry is vetted extensively by the administrators (our curation procedures are fully described in the Supplemental Information). While these ≈ 300 entries include those we consider to be essential for a quantitative understanding of human impacts on Earth, it is not an exhaustive list. This database will continue to grow and evolve as more data becomes publicly released, our understanding of the human-earth system improves, and members of the scientific community suggest values to be added.

To illustrate the structure of a database entry, let’s consider the most emblematic one: the atmospheric CO_2_ concentration as measured by the Mauna Loa Observatory (Figure 1). At the top of the entry we find the title and category (Fig. 1A-B). Primary categorization falls into one of five classes: “Land”, “Water”, “Energy”, “Flora & Fauna”, and “Atmospheric & Biogeochemical Cycles”. Of course, these categories are broad and entries can be associated with several categories. For this reason, each entry is also assigned a narrower “subcategory”, such as “Agriculture”, “Urbanization”, or “Carbon Dioxide.” While this categorization is not meant to be exhaustive, and many other schemes could be implemented, we found this organization allowed us to quickly browse and identify quantities of interest.

**Figure 1:**
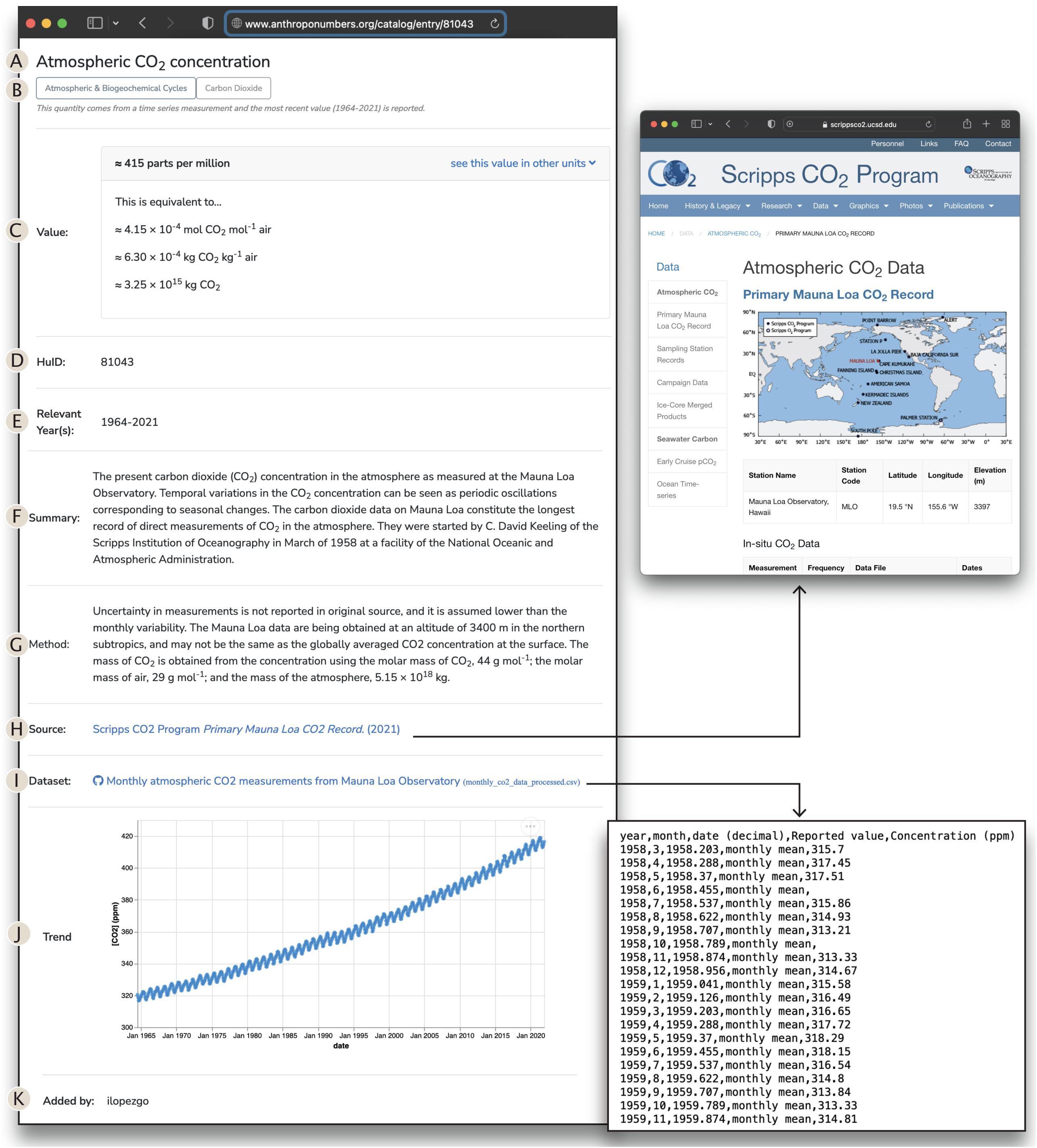
A representative entry in the Human Impacts Database. The entry page for HuID 81043 - “Atmospheric CO_2_ concentration” is diagrammed with important features highlighted. Each entry in the Human Impacts Database has a (A) name, (B) primary and secondary categorization, (C) the numerical value with other units when appropriate, (D) a 5-digit permanent numeric identifier, (E) years for which the measurement was determined, (F) a brief summary of the quantity, (G) the method of determination, (H) a link to the source data, and (I) a link to a processed version of the data saved as a .csv file. When possible, a time series of the data is presented. (K) Finally, each entry lists the username of the administrator who curated the quantity. Their contact information is available on the anthroponumbers.org “About” page.

Following the title and categorization, we report the measured atmospheric CO_2_ concentration. This corresponds to the most recent reported measurement, which is, as of writing, roughly 416 ppm in 2021 (parts per million, Figure 1C). Importantly, we report an approximate CO_2_ concentration rather than a precise one to many significant digits. While the most recent entry in the linked dataset (Fig. 1I) gives a monthly average value of 416.43 ppm for December of 2021, this value does not account for error in the measurement, fluctuations throughout December, or seasonal oscillations in atmospheric CO_2_. Therefore, we report a rounded value of 416 ppm. CO_2_ measurements are quite accurate, but other measurements and inferences recorded in the Human Impacts Database are less so. We therefore strive to give an assessment of the uncertainty for all values. This can be in the form of a confidence interval, as for HuID 11827, which reports a 90% confidence interval on the extent of sea level rise due to thermal expansion since 1900, or bounds on the value, as in HuID 44641, which reports a lower bound on the number of animal extinctions since 1500 CE. In addition to error assessment, we also aim to provide legible units for all entries. Though atmospheric CO_2_ is commonly reported in parts per million (ppm) units, we also report this value in other equivalent units, including the mole and mass fractions of CO_2_, and the total mass of CO_2_ in the atmosphere in kg CO_2_ (Fig. 1C). Whenever possible, entries will report values in multiple units to make quantities accessible to readers coming from diverse backgrounds.

Following the numerical value, we find the Human Impacts Database identifier (HuID, Figure 1D). The HuID is a randomly-generated five-digit number that serves as a permanent identifier of the entry. Because the HuID is permanent and static, it can be used for referencing. Rather than identifying a single value, we consider the HuID a pointer to a particular *entry*, so that HuID 81043 can be used to reference the atmospheric CO_2_ concentration in 2020 and 1980 (Fig. 1E). For example, to reference the present-day atmospheric CO_2_ concentration, one could report the value as “≈ 416 ppm (HuID 81043:2021)”. Additionally, since each entry comes from a single source, we may have more than one HuID reporting similar quantities, for example HuIDs 69674 and 72086 report recent measurements of the temperature of the upper ocean.

The “Summary” field (Figure 1F) gives a succinct description of the quantity and its relationship to “human impacts” broadly-construed, along with other pertinent information. This could include a more detailed definition of terms used in the quantity, such as the entry for “sea ice extent loss in March” which defines the term sea ice extent, or useful historical information about the measurement. In our example of atmospheric CO_2_ concentration, the summary explains that the measurement is made at the Mauna Loa observatory and points out the seasonal oscillations that are observed. The following “Method” field describes the method by which the quantity was measured, inferred, or estimated (Figure 1G). This field also provides an assessment of the uncertainty in the value, which may include a description of how confidence intervals were computed or a list of critical assumptions that were made to estimate missing data.

All fields through “Method” (Fig 1A-G) depend on manual curation and interpretation by database. The following two fields, “Source” and “Dataset” (Figure 1H-I) provide direct links to the primary source reference and the relevant data. Both of these fields are direct links (shown as insets in Figure 1). The “Source” can point either to the published scientific literature or the resource page of a governmental, industrial, or non-governmental organization data deposition URL. The “Dataset” field links directly to a CSV version of the datafile in our GitHub repository. As discussed in the Supplement, these data files have been converted into a “tidy-data” format^39^ by database administrators, which maximizes programmatic readability.

When possible, a graphical time-series of the data is also presented as an interactive plot (Fig. 1J). These plots enable users to quickly apprehend time-dependent trends in the data without downloading or processing the dataset. While not available for every entry, the majority of quantities we have curated in the Human Impacts Database contain measurements over time. The last field gives the username of the administrator who generated this entry (Figure 1K). Their affiliation and contact information is available on the database’s “About” page. We invite the reader to contact the administrators collectively - through our “Contact” page or directly through our personal emails as provided on the “About” page for questions, concerns, or suggestions.

While Figure 1 is a representative example, each quantity in the Human Impacts Database tells a different story. Easy centralized access to different entries allows users to learn about the magnitude of human impacts, and also study the interactions between different human activities, which, as we discuss in the next section, are deeply intertwined.

### Global Magnitudes

In Figure 2, we provide an array of quantities that we believe to be key in developing a “feeling for the numbers” associated with human impacts on the Earth system. All of the quantities in Figure 2 are drawn from entries in the database and grouped into the same categories used in the database: land, water, flora and fauna, atmosphere and biogeochemical cycles, and energy (see color scheme at the top of Fig. 2). Though the impacts considered necessarily constitute an incomplete description of human interaction with the planet, these numbers encompass many which are critically important, such as the volume of liquid water resulting from ice melt (Fig. 2B), the extent of urban and agricultural land use (Fig. 2H), global power consumption (Fig. 2N), and the heat uptake and subsequent warming of the ocean surface (Fig. 2S). In many cases, the raw numbers are astoundingly large and can therefore be difficult to fathom. Rather than reporting only bare “scientific” units, we present each quantity (when possible) in units that are relatable to a broad audience. For example, to give context for the annual global mass of CO_2_ emissions in kilograms, we note that this mass is equivalent to 2.5 two-tonne pickup trucks per person on the planet per year.

**Figure 2:**
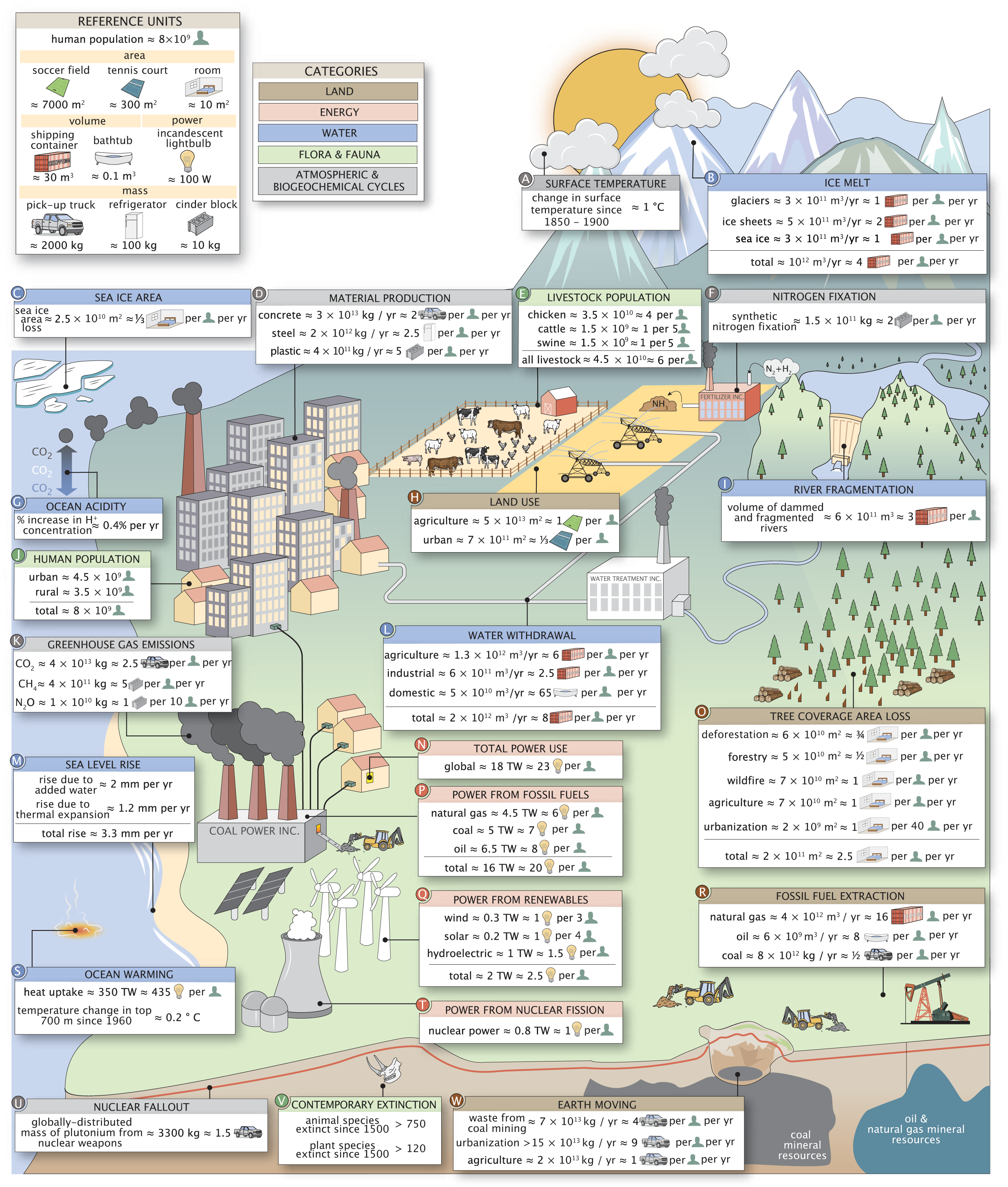
Human impacts on the planet and their relevant magnitudes. Relative units and the broad organizational categories are shown in the top-left panels. Source information and contextual comments for each subpanel are presented in the Supplemental Information.

Exploring these numbers reveals a number of intriguing quantities and relationships. For example, agriculture repeatedly appears as a major contributor to many human impacts. Agriculture dominates both global land (Fig. 2H) and water use (Fig. 2L), and accounts for approximately a third of global tree cover area loss (Fig. 2O). In addition, an enormous amount of nitrogen is synthetically fixed through the Haber-Bosch process to produce fertilizer (Fig. 2F), which is a major cause of emission of N_2_O (Fig. 2 K), which is a potent greenhouse gas. About 30 billion livestock are raised on agricultural lands (Fig. 2E), which, together with rice paddies, produce a majority of anthropogenic methane emissions (i.e. the greenhouse gas CH_4_, Fig. 2 K). On the other hand, urban land area accounts for a very small fraction of land area use (≈ 1%, Fig. 2H), and the expansion of cities and suburbs accounts for only ≈ 1% of global tree cover area loss (Fig. 2O). This is not to say, however, that urban centers are negligible in their global impacts. As urban and suburban areas currently house more than half of the global human population (Fig. 2J), many human impacts are linked to industries that directly or indirectly support urban populations’ demand for food, housing, travel, electronics and other goods. For example, the pursuit of urbanization is the dominating factor in the mass of earth moved on an annual basis (Fig. 2W).

Collectively, the ≈ 8 billion humans on Earth (Fig. 2J) consume nearly 20 TW of power, equivalent to 23 one hundred Watt light bulbs per person (Figure 2N). Around 80% of this energy derives from the combustion of fossil fuels (Fig. 2P). This results in a tremendous mass of CO_2_ being emitted annually (Fig. 2 K) of which only ≈ ½ remains in the atmosphere (HuID 70632). A sizable portion of the emissions are absorbed by the oceans (HuID 99089), leading to a steady increase in ocean acidity (Fig. 2G) and posing risks to marine ecosystems^40^. Furthermore, increasing average global temperatures primarily caused by greenhouse gas emissions contribute to sea level rise not only in the form of added water from melting ice (Fig. 2B and M) but also due to thermal expansion of ocean water (Fig. 2M), which accounts for ≈ 30% of observed sea level rise^41^. These are just a few ways in which one can traverse the impacts illustrated in Figure 2, revealing the remarkable extent to which these impacts are interconnected. We encourage the reader to explore this figure in a similar manner, blazing their own trail through the values.

### Regional Distribution

While Figure 2 presents the magnitude of human impacts at a global scale, it is important to recognize that these impacts — both their origins and repercussions — are variable across the globe. That is, different societies vary in their preferences for food (e.g. Americans consume relatively little fish) and modes of living (e.g. apartments vs houses), have different levels of economic development (e.g. Canada as compared to Malaysia), rely on different natural resources to build infrastructure (e.g. wood vs concrete) and generate power (e.g. nuclear vs coal), and promote different extractive or polluting industries (e.g. lithium mining vs palm oil farming). Some of these regional differences are evident in Figure 3, which summarizes regional breakdowns of several drivers of global human impacts, e.g. livestock populations and greenhouse gas emissions.

**Figure 3:**
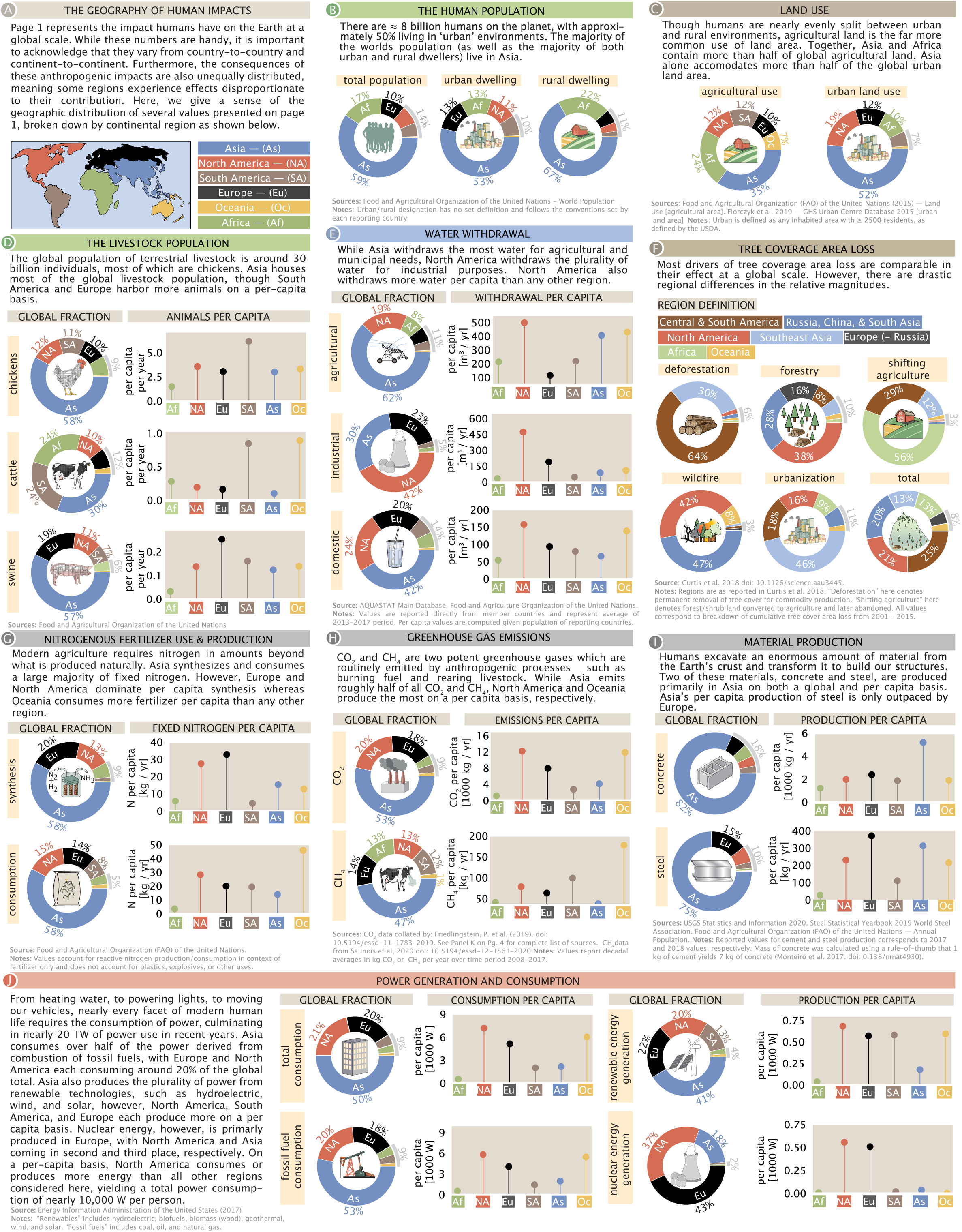
Regional distribution of anthropogenic effects. Several quantities from Figure 2 were selected and the relative magnitudes were broken down by subcontinental area (A). Donut charts in all sections show the relative contributions of each quantity by region. Ball-and-stick plots show the per capita breakdown of each quantity across geographic regions. All data for global and per-capita breakdowns correspond to the latest year for which data were available. The regional breakdown for deforestation uses the regional convention as reported in the source data^45^.

Just as impactful human activities like coal power generation and swine farming are more common in some regions than others (Fig 2.), likewise the impacts of human activities affect some regions more than others^42^. Figure 3 displays a coarse regional breakdown of the numbers from Figure 2 for which regional distributions could be determined from the literature. The region definitions used in Figure 3 are similar to the definitions set forth by the Food and Agricultural Organization (FAO) of the United Nations, assigning the semi-continental regions of North America, South America, Africa, Europe, Asia, and Oceania. Here, we specify both the total contribution of each region and the per capita value given the population of that region as of the year(s) in which the quantity was measured.

Much as in the case of our Figure 2, interesting details emerge naturally from Figure 3. For example, Asia dominates global agricultural water withdrawal (excluding natural watering via rainfall), using about 62% of the total, while North America takes the lead in industrial water withdrawal. Interestingly, on a per-capita basis North America withdraws the most water for all uses: agricultural, industrial, and domestic.

North America also emits more CO_2_ per capita than any other region, with Oceania and Europe coming second and third, respectively. This disparity can be partially understood by considering the regional distribution of fossil fuel consumption, the dominant source of CO_2_ emissions (Fig. 3J). While Asia consumes more than half of total fossil fuel energy, per capita consumption is markedly lower than in North America, Europe, and Oceania (Fig. 3J). Interestingly, the story is different when it comes to methane. Oceania and South America are the largest emitters of anthropogenic CH_4_, mainly due to a standing population of cattle that rivals that of humans in those regions (Fig. 3D) and produces this potent greenhouse gas through enteric fermentation^33^. Regional disparities are also apparent in the means of energy production. While consuming only 4% of total power, South America generates about 14% of renewable energy. Nuclear power generation, on the other hand, is dominated by North America and Europe, while Oceania, which has a single research-grade nuclear reactor, generates nearly zero nuclear energy.

Investigating the causes of forest loss by geographic region likewise highlights interesting differences. At a global level, all drivers of forest loss are comparable in magnitude except for urbanization, which accounts for ≈ 1% of total annual tree cover area loss (Fig. 2O). Despite comparable magnitudes, different drivers of forest loss have different long-term consequences^30^. Forest loss due to wildfires and forestry often result in regrowth, while commodity-driven harvesting and urbanization tend to be drivers of long-lasting deforestation^43,44^. Central and South America account for about 65% of commodity-driven deforestation (meaning, clearcutting and human-induced fires with no substantial regrowth of tree cover), whereas a majority of forest loss due to shifting agriculture occurs in Africa (where regrowth does occur). Together, wildfires in North America, Russia, China, and South Asia make up nearly 90% of losses due to fire^45^. While urbanization is the smallest driver of tree cover loss globally, it can still have strong impacts at the regional level, perturbing local ecosystems and biodiversity^46,47^.

### Timeseries

When available, the Human Impacts Database includes time series data for each quantity. Just as the regional distributions in impactful human activities help us understand differences between societies and regions, studying the history of these activities highlights recent technological and economic developments that intensify or reduce their impacts. When considering the history of human impacts on the Earth, it is natural to start by considering the growth of the human population over time. As shown in Figure 4, the global human population grew nearly continually over the past 80 years, with the current population nearing 8 billion. Historically, most of the global human population lived in rural areas (about 70% as of 1950, HuID 93995). Recent decades have been marked by a substantial shift in how humans live globally, with around half of the human population now living in urban or suburban settings (≈ 55%, HuID 93995).

**Figure 4:**
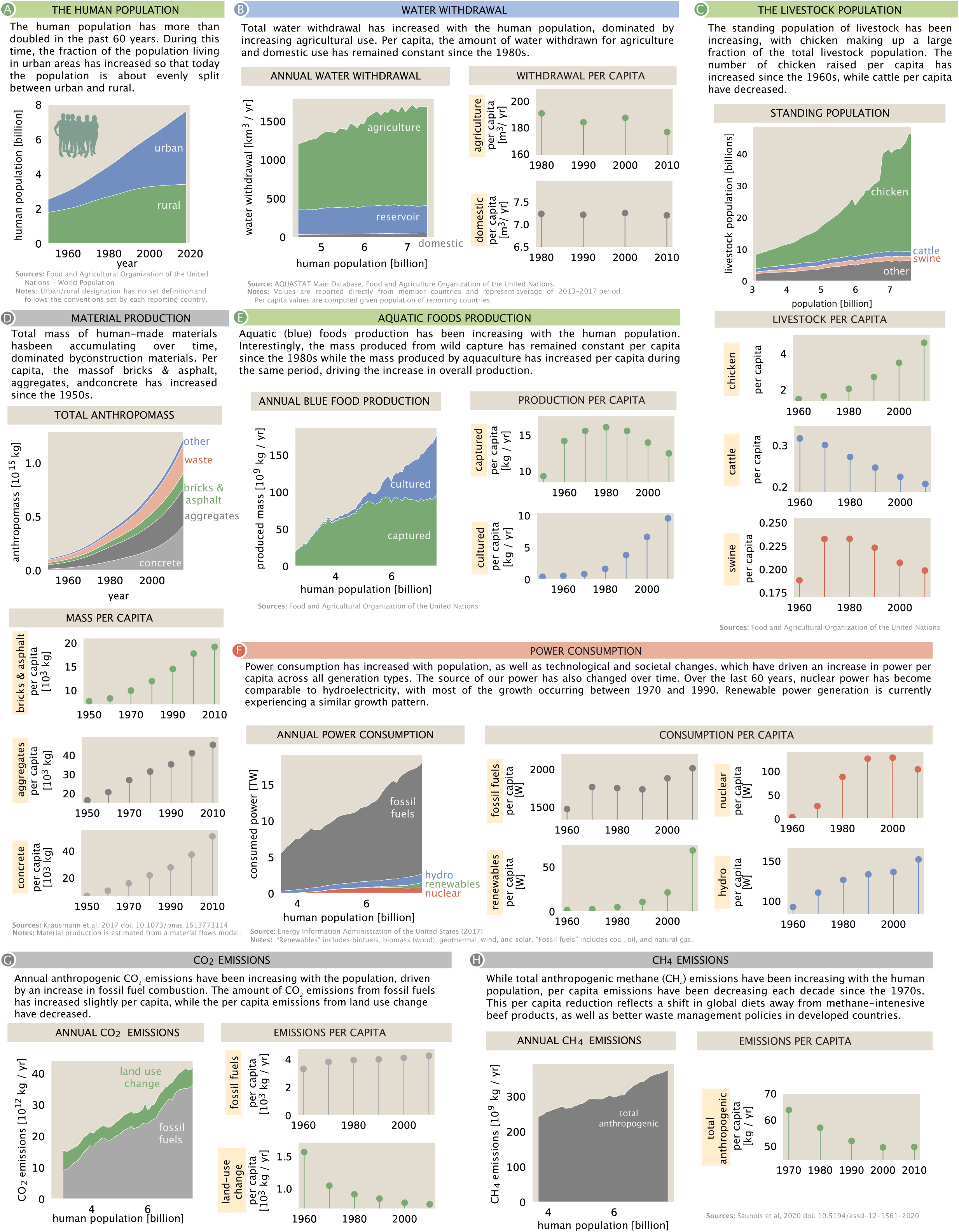
Temporal dynamics of key human impacts. Several quantities from Figure 2 were selected and the magnitudes were plotted either as a function of time (for cumulative quantities such as anthropomass) or human population (A). Ball-and-stick plots show the per capita breakdown as decadal averages to give a more reflective view of cultural and technological shifts than year-to-year variation.

Given the growth of the human population, it’s reasonable to consider that human population may be the most natural scale to measure human impacts^48^. To assess this possibility, we plot per-capita impacts over several decades (Figure 4). If impacts are growing in direct proportion with the human population, per capita impacts would be constant over time. Indeed, this is roughly true for per-capita water withdrawals over the last 40 years (Fig. 4B). Deviations from proportionality may indicate important changes in human activities. For example, in recent decades per-capita chicken populations grew by nearly two-fold while per-capita cattle populations shrunk by roughly 25%, reflecting a modest transition away from beef and towards chicken as a source of animal meat in global diets (HuIDs 40696 and 79776).

One very visible impact accompanying the shift of the human population to urban environments is the increase in production of anthropogenic mass -- materials such as concrete, steel, lumber, and plastics used to build roads, buildings, machines, packaging and other useful human-made items. Since these materials are degraded very slowly, anthropogenic mass has been accumulating over time. In addition, the mass of concrete, aggregates like asphalt, and bricks per capita has been increasing since the 1950s (Fig. 4D). Concrete, in particular, has increased from less than 10 tons per person in the 1950s to almost 30 tons per person in the 2010s. This increase in per capita anthropogenic mass means that the increase in production of these materials is outpacing the growth of the human population.

These material production trends have been enabled, in part, by a sustained increase in power generation. As evident from Figure 4, total power consumption has been increasing roughly proportionally with the human population. Per capita consumption has also increased across all generation types, including fossil fuels, hydropower, nuclear, and renewables. The growth among nuclear and renewables has been especially dramatic, and nuclear power now roughly equals hydropower production. Production of crops, aquaculture, and populations of livestock are all likewise correlated with growth in the human population (Fig. 4C and E). The total number of livestock has increased with the human population, primarily due to increasing chicken populations as discussed above. The dominant means of aquatic food production has also shifted over this time: until roughly 1980, nearly all seafood was captured wild, but since then aquaculture has grown to account for roughly ½ of aquatic food production (HuID 61233, Fig. 4E).

Turning our focus to greenhouse gases, we see that annual anthropogenic CO_2_ emissions have been increasing with the population (Fig. 4G). Burning of fossil fuels is the dominant contributor to anthropogenic emissions, and has increased slightly on a per capita basis over the past 60 years. In contrast, as the pace of global deforestation has slowed^49,50^ emissions of CO_2_ due to land-use change have decreased per capita. These two trends roughly neutralize each other, leading to little overall change in CO_2_ emissions per capita since the 1960s. Akin to CO_2_ emissions due to land-use change, CH_4_ emissions show a sublinear trend with human population, partially due to a decline in ruminant livestock per capita (Fig. 4C, H).

## Discussion

Quantitative literacy is necessary for understanding in nearly all branches of science. As our collective knowledge of anthropogenic impacts expands, it has become challenging to sift through the literature to collect specific numbers useful for both calculation and communication. We have attempted to reduce this barrier to entry on several fronts. We have canvassed the scientific literature, governmental and international reports to assemble a broad, quantitative picture of how human activities have impacted Earth’s atmosphere, oceans, rivers, lands, biota, chemistry, and geology. In doing so, we have created an online, searchable database housing an array of quantities and data that describe different facets of the human-Earth interface. Beyond the database, we have assembled these data into a comprehensive snapshot, released alongside this writing as a standalone graphical document (Supplemental File 1), with all underlying data, associated uncertainties and referencing housed in the Human Impacts Database. While necessarily incomplete, these resources provide a broad view of the ways in which human activities are impacting the Earth on multiple fronts.

One insight that emerges from considering these diverse human activities together is that they are deeply intertwined and driven by a small number of pivotal factors: the size of the human population, the composition of our diets, and our demand for materials and energy to build and power our increasingly complex and mechanized societies. Understanding the scale of human agriculture, water and power usage provides a framework for understanding most of the numerical gallery presented in Figure 2. Perhaps unsurprisingly, we find that feeding the growing human population is a major driver of a large swath of human impacts on earth, dominating global land (Figure 2H, HuID: 29582) and water use (Figure 2L, HuIDs: 84545, 43593, 95345), as well as significantly contributing to tree cover loss (Figure 2O, HuID: 24388), earth moving (Figure 2W, HuID: 19415, 41496), and anthropogenic nitrogen fixation (Figure 2F, HuID: 60580, 61614), to name a few such examples.

It is common in this setting to argue that the bewildering breadth and scale of human impacts should motivate some specific remediation at the global or local scale. We, instead, take a more modest “just the facts” approach. The numbers presented here show that human activities affect our planet to a large degree in many different and incommensurate ways, but they do not provide a roadmap for the future. Rather, we contend that any plans for the future should be made in the light of a comprehensive and quantitative understanding of the interconnected ways in which human activities impact the Earth system globally (Figure 2), locally (Figure 3), and temporally (Figure 4). Achieving such an understanding will require synthesis of broad literature across many disciplines. While the quantities we have chosen to explore are certainly not exhaustive, they represent some of the key axes which frequently drive scientific and public discourse and shape policy across the globe.

Earth is the only habitable planet we know of, so it is crucial to understand how we got here and where we are going. That is, how (and why) have human impacts changed over time? How are they expected to change in the future? For every aspect of human entanglement with the Earth system – from water use to land use, greenhouse gas emissions, mining of precious minerals, and so on – there are excellent studies measuring impacts and predicting their future trajectories. Of particular note are the data-rich and explanatory reports from the Intergovernmental Panel on Climate Change^51,52^ and the efforts towards defining “Planetary Boundaries”^53^. We hope that the Human Impacts Database and the associated resources with this work will aim another lens on the human-Earth system, one that engages the scientific community ultimately helping humanity coexist stably with the only planet we have.

## Supporting information

Graphical Snapshot

