## Supplemental Text for "Anthroponumbers.org: A Quantitative Database Of Human Impacts on Planet Earth"

**Supplementary Information for “Anthroponumbers.org: A Quantitative Database Of Human Impacts on Planet Earth”**

### Selection, Validation, and Curation of Values

The breadth of measurements of human impacts on the planet is enormous, covering a wide array of disciplines and methods. While this is a boon for science, this imposes a very important burden – any value we care to enter into the Human Impacts database must be carefully examined and deemed credible and appropriate for the database. While we certainly acknowledge we are not domain experts in *all* of these fields, the members of the administrative team span a broad range of backgrounds, and are all quantitative scientists who both deeply value the utility of quantitative measurements and have the domain expertise to assess whether the reported values make sense and are determined with trustworthy methods. In this section, we briefly outline the general procedure undertaken before a value is entered into the Human Impacts Database.

### Identifying a Potential Entry

Our first action is identifying a value or set of values and determining whether they are pertinent to Human Impacts. We take a broad definition of “Human Impacts”, but enforce that any value must be either (i) a direct result of anthropogenic action, (ii) contributed to by anthropogenic activities, or (iii) is directly relevant to human consumption and/or production. Most importantly, any candidate entry must reflect an impact on some natural process. For example, a value quantifying the standing population of all livestock on Earth would fall under criterion (i) making it an appropriate candidate entry. As a counter example, the fraction of a country’s GDP resulting from fossil fuel export would not be considered as a candidate value as it describes an economic impact rather than an impact on a natural process. Of course, rigid lines cannot always be drawn and inclusion of a value is ultimately at the discretion of the administration team.

### Vetting a Potential Entry

Next, we determine if the quantity is scientifically valid and appropriate. This not only includes the precise value of the quantity, but the reliability of the source and the methods of measurement.

In general, we consider data from large, international efforts such as the Food and Agriculture Organization of the UN (FAO) or the Intergovernmental Panel on Climate Change (IPCC) to be highly reliable sources of information. We take these sources to be reliable as they clearly report the methods of their measurements or meta-analyses, emphasizing where assumptions and approximations have been made. Furthermore, given the internationality of its contributors and the deep well of scientists they consult and employ, we find that the FAO and IPCC are largely free of bias as they have little stake in reporting overly-rosy or negative results. For this reason, we are less likely to include values from industry reports, which have potential conflicts of interest. Whenever industry reports are used, we try to find multiple sources for that particular value to place it in context. For example, we extensively use the BP Annual Statistical Report on Energy in the human impacts database. As BP is a private company with financial interests in reporting global energy use, we compare these values with those from the US Energy Information Administration (EIA) and the International Energy Agency (IEA) to judge their consistency.

We draw a large number of the values in the Human Impacts Database from peer-reviewed scientific reports. For these data sources we thoroughly examine the reported methods used to determine the value. If details regarding the method are not clearly reported (e.g. the value “was fitted” without explaining the fitting procedure), we are strongly inclined to not trust that particular source. Furthermore, if the method is not stated or the code/data are available under only “reasonable request”, the value is not considered as appropriate. When possible, we also compare the reported value to other measurements and check if the source explains any discrepancy between their measurement and others. In many cases, however, there are not multiple reported values for a given quantity. In these cases, we assess the trustworthiness of the reported value and reach out to domain experts as needed. With rare exceptions, we do not factor the publishing journal in assessing the veracity of a value.

Once a value is entered into the database, we label it with a primary and secondary category. Human impacts are inherently connected by webs of interactions and often affect multiple subsystems within the Earth system. Meanwhile, most human impacts can be categorized according to the systems with which they interact most strongly. While incomplete, these category labels are meant to give users an impression of the subsystems that are most strongly influenced by or related to the value.  Users are able to filter the database by these categories and subcategories.

**Considering Uncertainty**

While the numeric value of a candidate quantity is an important factor we consider, so too is the reported uncertainty. Many scientific reports will give an assessment of uncertainty, either at the statistical, measurement, or systematic level. The clarity of the presented uncertainty analysis is critical in our determination of whether a candidate value should be entered in the database. While scientific reports often address the uncertainty, this is rarely reported in governmental and industry reports. Many numbers from governmental or intergovernmental bodies come from surveys and are thus self-reported by countries, adding some uncertainty to the data and requiring some level of interpolation from the reporting agency. These numbers are still considered, though we are cognizant of the number of significant digits that are reported. Often, we report these numbers as approximate, representing the uncertainty with the data. In all cases, we state a concise yet sufficiently detailed description of the method and quantification of uncertainty in the “method” field of an HuID entry.

**Considering Data Use Protections**

As we do not directly generate the data presented in this work, we are very careful to ensure that the data we add to the database follows all legal requirements. All data presented in the database must be explicitly stated to be under a generally permissive license such as a Creative Commons Attribution license (CC-BY). Data sources which reserve all rights to their data are not included in the database in any form. While we ensure that we have the legal right to share these data, we strongly implore the users of the Human Impacts Database to directly cite the original data source alongside the database if a value or entry is used in a later publication.

**Continued Curation and Maintenance of the Database**

Unlike similar databases in chemistry and biology (such as BioNumbers or the CRC Handbook of Chemistry and Physics), the Human Impacts Database faces a unique maintenance challenge as the values it houses will undoubtedly change with time as will our understanding of the facets of the Earth system that are impacted by human activities. This means that a concerted effort to keep the values in the database up to date, within reason, is needed. In this section, we outline steps we have taken to ensure that the database can be properly maintained and be useful for many years to come.

*Composition of the Administrative Team*

The primary authors on this work (GC, RAB, AIF, ILG, NSS, and MK) are the primary members of the administrative team of the Human Impacts Database. All of these authors are practicing research scientists working at the interface of biology, chemistry, physics, and earth science. As a result, this database will be an invaluable resource for our specific research objectives, imposing a self-interest in keeping the entries up to date. All members of the administrative team frequently read primary scientific literature covering these topics, meaning that critical new values or updates to extant entries can be reliably found. Furthermore, the majority of the administrative team intend to enter into leadership positions in academic and industry contexts, allowing us to mentor and train more administrators with different domain expertises. As this database is primarily a scientific tool, we believe our specific yet diverse training well prepares us as careful curators of the database. Furthermore, all authors are well-versed in computational methods with some administrators having expertise in web development technologies. This added expertise helps ensure that the database will reliably operate at both the front and backend levels. In addition, the two PIs who have led this work, RP and RM, have support from the Resnick Sustainability Center at Caltech and the Weizmann Institute to continue work on this project.

Many of the sources behind the HuID entries are updated on a regular basis, but updates may not be immediately updated on the database itself. For example, the FAO routinely updates their data as new data arrive or corrections/improvements to previously reported data are released. The frequent nature of these releases precludes a mirror reflection of these values in the Human Impacts Database. For continually updating sources, we update these values at an annual basis within the third quarter of the calendar year. Other sources, such as the BP statistical report on energy and IPCC reports, also typically release updates around this time. For values that are more frequently updated (such as the atmospheric $CO_2$ concentration, which is updated on a near-daily basis), we update these values semiannually coinciding with the spring and fall of the calendar year.

While the administration team is diverse in their scientific interests and expertise, it is unreasonable to believe that our collective knowledge is all-encompassing of Human Impacts. There will invariably be important values that we are unaware of that should be included in the database. To this end, we have developed a community-feedback system into the database (https://anthroponumbers.org/catalog/contact) where the general public can submit recommendations for new values or updates and/or corrections to extant values in the database. Whenever feedback is submitted, the administrative team is notified, preventing important feedback from being cast into the void. Furthermore, contact information is provided for each administrative member (https://anthroponumbers.org/catalog/about) if a user wishes to contact us individually.

As the curation procedures enumerated in the preceding sections are laborious and require a level of comfort in digesting scientific methods and data, we have opted to not open core maintenance privileges to the general public. However, all values housed within the database are also housed within a public GitHub repository (https://github.com/rpgroup-pboc/human_impacts) where we enthusiastically encourage forking of the repository and submission of new issues and pull requests. The issues and pull requests are also monitored by the administrative team.

### References and Explanations For Values Reported in Figure 1

In this section, we report our extensive and detailed referencing for each and every quantity reported in the subpanels of Figure 1 of the main text. As described in the Materials & Methods, each value comes from the manual curation of a piece of scientific, industrial, governmental, or non-governmental organization reports, articles, or databases. Each value listed here contains information about the original source, the method used to obtain the value, as well as accession identification numbers for the Human Impacts Database (https://anthroponumbers.org), listed as HuIDs.

For each value, we attempt to provide an assessment of the uncertainty. For some values, this corresponds to the uncertainty in the measurement or inference as stated in the source material. In cases where a direct assessment of the uncertainty was not clearly presented, we sought other reported values for the same quantity from different data sources to present a range of the values.

For others, this uncertainty represents the upper- and lower-bounds of the measurement or estimation.

Each value reported here is prefixed with a symbol representing our confidence in the value. Cases in which an equality (=) symbol is used represents that a measure of the uncertainty is reported in the original data source or represents a range of values from different sources that are tightly constrained (with 2 significant digits). An approximation symbol (≈) indicates values that we are confident in to within a factor of a few. In some cases, an approximation symbol (≈) represents a range where the values from different sources differ within three significant digits. In these cases, the ranges are presented as well. Finally, in some cases only a lower-bound for the quantity was able to be determined. These values are indicated by the use of an inequality symbol (>).

### A. Surface Warming

*Surface temperature change from the 1850-1900 average ≈ 1.0 - 1.3 ℃ (HuID: 79598, 76539, 12147)*

**Data Source(s):** HadCRUT.4.6 (Morice et al., 2012, DOI: 10.1029/2011JD017187), GISTEMP v4 (GISTEMP Team, 2020: GISS Surface Temperature Analysis (GISTEMP), version 4. NASA Goddard Institute for Space Studies. Dataset accessed 2020-12-17 at https://data.giss.nasa.gov/gistemp/ & Lenssen et al., 2019, DOI: 10.1029/2018JD029522) and NOAAGlobalTemp v5 (Huang et al, 2020, DOI: 10.1175/JCLI-D-19-0395.1) datasets.

**Notes:** The global mean surface temperature captures near-surface air temperature over the planet's land and ocean surface. The value reported represents the spread of three estimates and their 95% confidence intervals for the year 2019. Since data for the period 1850-1880 are missing in GISTEMP v4 and NOAAGlobalTemp v5, data are centered by setting the 1880-1900 mean of all datasets to the HadCRUT.4.6 mean over the same period.

### B. Annual Ice Melt

*Glaciers = $(3.0 \pm 1.2) \times 10^{11}$ $m^3$ / yr (HuID: 32459)*

**Data Sources:** Intergovernmental Panel on Climate Change (IPCC) 2019 Special Report on the Ocean and Cryosphere in a Changing Climate. Table 2.A.1 on pp. 199-202.

**Notes:** Value corresponds to the trend of annual glacial ice volume loss (reported as ice mass loss) from major glacierized regions (2006-2015) based on aggregation of observation methods (original data source: Zemp et al. 2019, DOI:10.1038/s41586-019-1071-0) with satellite gravimetric observations (original data source: Wouters et al. 2019, DOI:10.3389/feart.2019.00096). Ice volume loss was calculated from ice mass loss assuming a standard pure ice density of 920 kg / $m^3$. Uncertainty represents a 95% confidence interval calculated from standard error propagation of the 95% confidence intervals reported in the original sources assuming them to be independent.

*Ice sheets = $(4.6 \pm 0.4) \times 10^{11}$ $m^3$ / yr (HuIDs: 95798; 93137)*

**Data Source(s):** D. N. Wiese et al. 2019 JPL GRACE and GRACE-FO Mascon Ocean, Ice, and Hydrology Equivalent HDR Water Height RL06M CRI Filtered Version 2.0, Ver. 2.0, PO.DAAC, CA, USA. Dataset accessed [2022-Feb-09]. DOI: 10.5067/TEM- SC-3MJ62

**Notes:** Value corresponds to the trends of combined annual ice volume loss (reported as ice mass loss) from the Greenland and Antarctic Ice Sheets (2002-2021) measured by satellite gravimetry. Ice volume loss was calculated from ice mass loss assuming a standard pure ice density of 920 kg / $m^3$. Uncertainty represents one standard deviation and considers only propagation of monthly uncertainties in measurement.

*Arctic sea ice = (3.0 ± 1.0) × $10^{11}$ $m^3$ / yr (HuID: 89520)*

**Data Source(s):** PIOMAS Arctic Sea Ice Volume Reanalysis, Figure 1 of webpage as of January 31, 2022. Original method source: Schweiger et al. 2011, DOI:10.1029/2011JC007084

**Notes:** Value reported corresponds to the trend of annual volume loss from Arctic sea ice (1979-2022). The uncertainty in the trend represents the range in trends calculated from three ice volume determination methods.

### C. Sea Ice Area

*Extent of loss at yearly maximum cover (September) ≈ 4.8 × $10^{10}$ $m^2$ / yr (HuID: 66277)*

*Extent loss at yearly minimum cover (March) ≈ 0.4 × $10^{10}$ $m^2$ / yr (HuID: 66277)*

*Average annual extent loss = 2.5 × $10^{10}$ $m^2$ / yr (HuID: 66277)*

**Data Source(s):** Fetterer et al. 2017, updated daily. Sea Ice Index, Version 3, Boulder, Colorado USA. NSIDC: National Snow and Ice Data Center, DOI:10.7265/N5K072F8, [Accessed 2022-Feb-16].

**Notes:** Sea ice area is calculated by multiplying the percentage of sea ice in each pixel by pixel area and taking the integral sum of these products. Annual value corresponds to the linear trend of annual extent loss calculated by averaging over every month in a given year (2.45 × $10^{10}$ $m^2$ / yr HuID: 66277). The minimum cover area loss corresponds to the linear trend of Arctic sea ice area in September from 1979-2021 and the maximum cover area loss corresponds to the linear trend of sea ice area in March from 1979-2021. The Antarctic sea ice area trend is not shown because a significant long-term trend over the satellite observation period is not observed and short-term trends are not yet identifiable.

### D. Annual Material Production

*Concrete production ≈ (2 - 3) × $10^{13}$ kg / yr (HuID: 25488; 81346; 16995)*

**Data Source(s):** United States Geological Survey (USGS) National Minerals Information Center, Commodity Statistics and Information, Cement Statistics and Information. Miller et al. 2016, Table 1, DOI:10.1088/1748-9326/11/7/074029. Monteiro et al. 2017, DOI:10.1038/nmat4930. Krausmann et al. 2017, DOI:10.1073/pnas.1613773114

**Notes:** Concrete is formed when aggregate material is bonded together by hydrated cement. The USGS reports the mass of cement produced in 2019 as 4.1 × $10^{12}$ kg. As most cement is used to form concrete, cement production can be used to estimate concrete mass using a multiplicative conversion factor of 7 (Monteiro et al.). Miller et al. report that the cement, aggregate and water used in concrete in 2012 sum to 2.3 × $10^{13}$ kg. Krausmann et al. report an estimated value from 2010 based on a material input, stocks, and outputs model. The value is net annual addition to concrete stocks plus annual waste and recycling to estimate gross production of concrete.

*Steel production ≈ 1.9 × $10^{12}$ kg / yr (HuID: 51453; 44894; 85981)*

**Data Source(s):** United States Geological Survey (USGS) National Minerals Information Center, Commodity Statistics and Information, Iron and Steel Statistics and Information. World Steel Association, World Steel in Figures 2021, p. 7. Krausmann et al. 2017, DOI:10.1073/pnas.1613773114

**Notes:** Crude steel includes stainless steels, carbon steels, and other alloys. The USGS reports the mass of crude steel produced in 2019 as 1.860 × $10^{12}$ kg. The World Steel Association reports a production value of 1.874 × $10^{12}$ kg in 2019. Krausmann et al. report an estimated value from 2010 based on a material input, stocks, and outputs model. The value is net annual addition to steel stocks plus annual waste and recycling to estimate gross production of steel.

*Plastic production ≈ 4 × $10^{11}$ kg / yr (HuID: 97241; 25437)*

**Data Source(s):** Geyer et al. 2017, Table S1, DOI:10.1126/sciadv.1700782. Krausmann et al. 2017, DOI:10.1073/pnas.1613773114

**Notes:** Value represents the approximate sum total global production of plastic fibers and plastic resin during the calendar year of 2015. Comprehensive data about global plastic production is sorely lacking. Geyer et al. draw data from various industry groups to estimate total production of different polymers and additives. Some of the underlying data is not publicly available, and data from financially-interested parties is inherently suspect. Krausmann et al. report an estimated value from 2010 based on a material input, stocks, and outputs model. The value is net annual addition to stocks plus annual waste and end-of-life recycling to estimate gross production of plastics.

### E. Livestock Population

*Chicken standing population ≈ $3.5 \times 10^{10}$ (HuID: 94934)*
*Cattle standing population ≈ $1.5 \times 10^{9}$ (HuID: 92006)*
*Swine standing population ≈ $1.5 \times 10^{9}$ (HuID: 21368)*
*All livestock standing population ≈ $4.6 \times 10^{10}$ (HuID: 43599)*

**Data Source(s):** Food and Agriculture Organization (FAO) of the United Nations Statistical Database (2022) — Live Animals.

**Notes:** Counts correspond to the estimated standing populations in 2019. Values are reported directly by countries. The FAO uses non-governmental statistical sources to address uncertainty and missing (non-reported) data. Reported values are therefore approximations.

### F. Annual Synthetic Nitrogen Fixation

*Annual mass of synthetically fixed nitrogen ≈ $(1.4 - 1.5) \times 10^{11}$ kg N / yr (HuID: 60580; 61614)*

**Data Source(s):** United States Geological Survey (USGS) National Minerals Information Center, Commodity Statistics and Information, Nitrogen Statistics and Information. International Fertilizer Association (IFA) Statistical Database (2021) — Ammonia Production & Trade Tables by Region. Smith et al. 2020, DOI: 10.1039/c9ee02873k.

**Notes:** Ammonia ($NH_3$) produced globally is compiled by the USGS and IFA from major factories that report output. The USGS estimates the approximate mass of nitrogen in ammonia produced in 2019 as $1.42 \times 10^{11}$ kg N and the International Fertilizer Association reports a production value of $1.50 \times 10^{11}$ kg N in 2019. Nearly all of this mass is produced by the Haber-Bosch process (>96%, Smith et al. 2020). In the United States most of this mass is used for fertilizer, with the remainder being used to synthesize nitrogen-containing chemicals including explosives, plastics, and pharmaceuticals (≈ 88%, USGS Mineral Commodity Summaries 2020 – Nitrogen).

### G. Ocean Acidity

Surface ocean [H+] ≈ 0.2 parts per billion (HuID: *90472*)
Annual change in [H+] = *0.36 ± 0.03% (HuID: 19394)*

**Data Source(s):** Figures 1-2 of European Environment Agency report CLIM 043 (2020). Original data source of the report is "Global Mean Sea Water pH" from Copernicus Marine Environment Monitoring Service.

**Notes:** Reported value is calculated from the global average annual change in pH over years 1985-2018. The average oceanic surface pH was ≈ 8.057 in 2018 and decreases annually by ≈ 0.002 units, giving a change in [H+] of roughly $10^{-8.055} - 10^{-8.057} \approx 4 \times 10^{-11}$ mol/L or about 0.4% of the global average. [H+] is calculated as $10^{-pH} \approx 10^{-8}$ mol/L or 0.2 parts per billion (ppb), noting that $[H_2O] \approx 55$ mol/L. Uncertainty for annual change is the standard error of the mean.

### H. Land Use

*Agriculture ≈ 5 × $10^{13}$ $m^2$ (HuID: 29582)*
**Data Source(s):** Food and Agriculture Organization (FAO) of the United Nations Statistical Database (2020) — Land Use.
**Notes:** Agricultural land is defined as all land that is under agricultural management including pastures, meadows, permanent crops, temporary crops, land under fallow, and land under agricultural structures (such as barns). Reported value corresponds to 2017 estimates by the FAO.
*Urban ≈ (6 - 8) × $10^{11}$ $m^2$ (HuID: 41339; 39341)*
**Data Source(s):** Florczyk et al. 2019 (https://tinyurl.com/yyxxgtll) and Table 3 of Liu et al. 2018 DOI: 10.1016/j.rse.2018.02.055
**Notes:** Urban land area is determined from satellite imagery. An area is determined to be "urban" if the total population is greater than 5,000 and has a minimum population density of 300 people per $km^2$. Reported value gives the range of recent measurements of ≈ 6.5×$10^{11}$ $m^2$ (2015) and ≈ (7.5 ± 1.5) ×$10^{11}$ $m^2$ (2010) from Florczyk et al. 2019 and Liu et al. 2018, respectively.

### I. River Fragmentation

*Global fragmented river volume ≈ 6 × $10^{11}$ $m^3$ (HuID: 61661)*
**Data Source(s):** Grill et al. 2019 DOI: 10.1038/s41586-019-1111-9
**Notes:** Value corresponds to the water volume contained in rivers that fall below the connectivity threshold required to classify them as free-flowing. Value considers only rivers with upstream catchment areas greater than 10 $km^2$ or discharge volumes greater than 0.1 $m^3$ per second. The ratio of global river volume in disrupted rivers to free-flowing rivers is approximately 0.9. The exact value depends on the cutoff used to define a "free-flowing" river. We direct the reader to the source for thorough detail.

### J. Human Population

*Urban population ≈ 55% (HuID: 93995)*
*Global population ≈ 7.6 × $10^9$ people (HuID: 85255)*
**Data Source(s):** Food and Agricultural Organization (FAO) of the United Nations Report on Annual Population, 2019.
**Notes:** Value for total population in 2018 comes from a combination of direct population reports from country governments as well as inferences of underreported or missing data. The definition of "urban" differs between countries and the data does not distinguish between urban and suburban populations despite substantive differences between these land uses (Jones & Kammen 2013, DOI: 10.1021/es4034364). As explained by the United Nations population division, "When the definition used in the latest census was not the same as in previous censuses, the data were adjusted whenever possible so as to maintain consistency." Rural population is computed from this fraction along with the total human population, implying that the total population is composed only of "urban" and "rural" communities.

### K. Greenhouse Gas Emissions

*Anthropogenic $CO_2$ = (4.25 ± 0.33) × $10^{13}$ kg $CO_2$ / yr (HuID: 24789; 54608; 98043; 60670)*
**Data Source(s):** Table 6 of Friedlingstein et al. 2019, DOI: 10.5194/essd-11-1783-2019. Original data sources relevant to this study compiled in Friedlingstein et al.: 1) Gilfillan et al. https://energy.appstate.edu/CDIAC 2) Average of two bookkeeping models: Houghton and Nassikas 2017 DOI: 10.1002/2016GB005546; Hansis et

al. 2015 DOI: 10.1002/2014GB004997. 3) Dlugokencky and Tans, National Oceanic & Atmospheric Administration, Earth System Research Laboratory (NOAA/ESRL), https://www.esrl.noaa.gov/gmd/ccgg/trends/global.html, [Accessed 3-Nov-2019].

**Notes:** Value corresponds to total $CO_2$ emissions from fossil fuel combustion, industry (predominantly cement production), and land-use change during calendar year 2018. Emissions from land-use change are due to the burning or degradation of plant biomass. In 2018, roughly $1.88 \times 10^{13}$ kg $CO_2$ / yr accumulated in the atmosphere, reflecting the balance of emissions and $CO_2$ uptake by plants and oceans (Dlugokencky and Tans). Uncertainty corresponds to one standard deviation.

*Anthropogenic $CH_4$ = (3.4 - 3.9) × $10^{11}$ kg $CH_4$ / yr (HuID: 96837; 30725)*

**Data Source(s):** Table 3 of Saunois, et al. 2020. DOI: 10.5194/essd-12-1561-2020.

**Notes:** Value corresponds to 2008-2017 decadal average mass of $CH_4$ emissions from anthropogenic sources. Includes emissions from agriculture and landfill, fossil fuels, and burning of biomass and biofuels, but other inventories of anthropogenic methane emissions are also considered. Reported range represents the minimum and maximum estimated emissions from a combination of “bottom-up” and “top-down” models.

*Anthropogenic $N_2O$ = 1.1 (+0.6, -0.5) × $10^{10}$ kg $N_2O$ / yr (HuID: 44575)*

**Data Source(s):**Table 1 of Tian, H., et al. 2020. DOI: 10.1038/s41586-020-2780-0

**Notes:** Value corresponds to annualized $N_2O$ emissions from anthropogenic sources in the years 2007-2016. The value reported in the source is 7.3 [4.2, 11.4] Tg N / year. This is converted to a mass of $N_2O$ using the fact that N ≈ 14/22 of the mass of $N_2O$. Reported value is mean with the uncertainty bounds (+,-) representing the maximum and minimum values observed in the 2007-2016 time period.

### L. Water Withdrawal

*Agricultural = 1.3 × $10^{12}$ $m^3$ / year (HuID: 84545, 43593, 95345)*

*Industrial = 5.9 × $10^{11}$ $m^3$ / year (HuID: 27142)*

*Domestic = 5.4 × $10^{10}$ $m^3$ / year (HuID: 69424)*

*Total = (1.7 - 2.2) × $10^{12}$ $m^3$ / year (HuID: 27342, 68004)*

**Data Source(s):** Figure 1 of Qin et al. 2019. DOI: 10.1038/s41893-019-0294-2. AQUASTAT Main Database, Food and Agriculture Organization of the United Nations

**Notes:** “Agricultural” and “total” withdrawal include one value from Qin et al. (who reports “consumption”) and one value from the AQUASTAT database. Industrial water withdrawal is from AQUASTAT and domestic withdrawal value is from Qin et al. Values in AQUASTAT are self-reported by countries and have missing values from some countries, probably accounting for a few percent underreporting. All values represent water withdrawals. For agricultural and domestic, water withdrawal is assumed to be the same as water consumption, which is reported in Qin et al.

### M. Sea Level Rise

*Added water = 1.97 (+0.36, -0.34) mm / yr (HuID: 97108)*

*Thermal expansion = 1.19 (+0.25, -0.24) mm / yr (HuID: 97688)*

*Total observed sea-level rise = 3.35 (+0.47, -0.44) mm / yr (HuID: 81373)*

**Data Source(s):** Table 1 of Frederikse et al. 2020. DOI:10.1038/s41586-020-2591-3.

**Notes:** Values correspond to the average global sea level rise for the years 1993 - 2018. “Added water” (barystatic) change includes effects from meltwater from glaciers and ice sheets, added mass from sea-ice discharge, and changes in the amount of terrestrial water storage. Thermal expansion accounts for the volume change of water with increasing temperature. Values for “thermal expansion” and “added water” come from direct observations of ocean temperature and gravimetry/altimetry, respectively. Total sea level rise is the observed value using a combination of measurement methods. “Other sources” reported in Figure 1 accounts for observed residual sea level rise not attributed to a source in the model. Values in brackets correspond to the upper and lower bounds of the 90% confidence interval.

### N. Total Power Use

*Global power use ≈ 19 - 20 TW (HuID: 31373; 85317)*

**Data Source(s):** bp Statistical Review of World Energy, 2020; U.S. Energy Information Administration, 2020.

**Notes:** Value represents the sum of total primary energy consumed from oil, natural gas, coal, and nuclear energy and electricity generated by hydroelectric and other renewables. Value is calculated using annual primary energy consumption as reported in data sources assuming uniform use throughout a year, yielding ≈ 19 - 20 TW.

### O. Tree Coverage Area Loss

*Commodity-driven deforestation = (5.7 ± 1.1) × $10^{10}$ $m^2$ / yr (HuID: 96098)*
*Forestry = (5.4 ± 0.8) × $10^{10}$ $m^2$ / yr (HuID: 38352)*
*Urbanization = (2 ± 1) × $10^9$ $m^2$ / yr (HuID: 19429)*
*Shifting agriculture = (7.5 ± 0.9) × $10^{10}$ $m^2$ / yr (HuID: 24388)*
*Wildfire = (7.2 ± 1.3) × $10^{10}$ $m^2$ / yr (HuID: 92221)*
*Total tree cover area loss ≈ 2 × $10^{11}$ $m^2$ / yr (HuID: 78576)*

**Data Source(s):** Table 1 of Curtis et al. 2018 DOI:10.1126/science.aau3445. Hansen et al. 2013 DOI:10.1126/science.1244693. Global Forest Watch, 2020. Reported values in source correspond to total loss from 2001 - 2015. Values given are averages over this 15 year window.

**Notes:** Commodity-driven deforestation is “long-term, permanent, conversion of forest and shrubland to a non-forest land use such as agriculture, mining, or energy infrastructure.” Forestry is defined as large-scale operations occurring within managed forests and tree plantations with evidence of forest regrowth in subsequent years. Urbanization converts forest and shrubland for the expansion and intensification of existing urban centers. Disruption due to “shifting agriculture” is defined as “small- to medium-scale forest and shrubland conversion for agriculture that is later abandoned and followed by subsequent forest regrowth”. Disruption due to wildfire is “large-scale forest loss resulting from the burning of forest vegetation with no visible human conversion or agricultural activity afterward.” Uncertainty corresponds to the reported 95% confidence interval. Uncertainty is approximate for “urbanization” as the source reports an ambiguous error of “± <1%.”

### P. Power From Fossil Fuels

*Natural gas = 4.5 - 4.9 TW (HuID: 49947; 86175)*
*Oil = 6.1 - 6.6 TW (HuID: 42121; 39756)*
*Coal = 5.0 - 5.6 TW (HuID: 10400; 60490)*
*Total = 16 - 17.0 TW (HuID: 29470; 29109 )*

**Data Source(s):** bp Statistical Review of World Energy, 2020. U.S. Energy Information Administration, 2022.

**Notes:** Values are self-reported by countries. All values from bp Statistical Review and EIA correspond to 2019.. Reported TW values are computed from primary energy units (e.g. kg coal) assuming uniform use throughout the year. Oil volume includes crude oil, shale oil, oil sands, condensates, and natural gas liquids separate from specific natural gas mining. Natural gas value excludes gas flared or recycled and includes natural gas produced for gas-to-liquids transformation. Coal value includes 2019 value exclusively for solid commercial fuels such as bituminous coal and anthracite, lignite and subbituminous coal, and other solid fuels. This includes coal used directly in power production as well as coal used in coal-to-liquids and coal-to-gas transformations.

### Q. Power From Renewable Resources

*Wind = 0.36 - 0.43 TW (HuID: 30581, 85919)*
*Solar = 0.18 - 0.21 TW (HuID: 99885, 58303)*
*Hydroelectric = 1.2 - 1.3 TW (HuID: 15765, 50558)*
*Total = 1.9 - 2.1 TW (HuID: 74571, 20246)*
**Data Source(s):** bp Statistical Review of World Energy, 2020. U.S. Energy Information Administration, 2022.
**Notes:** Reported values correspond to estimates for the 2019 calendar year. Renewable resources are defined as wind, geothermal, solar, biomass and waste. Hydroelectric, while presented here, is not defined as a renewable in the BP dataset. All values are reported as input-equivalent energy, meaning the input energy that would have been required if the power was produced by fossil fuels. BP reports that fossil fuel efficiency used to make this conversion was about 40% in 2017.

### R. Fossil Fuel Extraction

*Natural gas volume = (3.9 - 4.0) × $10^{12}$ $m^3$ / yr (HuID: 11468; 20532)*
*Oil volume = (5.5 - 5.8) × $10^9$ $m^3$ / yr (HuID: 66789; 97719)*
*Coal mass = (7.8 - 8.1) × $10^{12}$ kg / yr (HuID: 78435; 48928)*
**Data Source(s):** bp Statistical Review of World Energy, 2020. U.S. Energy Information Administration (EIA), 2022.
**Notes:** Oil volume includes crude oil, shale oil, oil sands, condensates, and natural gas liquids separate from specific natural gas mining. Natural gas value excludes gas flared or recycled and includes natural gas produced for gas-to-liquids transformation. Coal value includes solid commercial fuels such as bituminous coal, anthracite, lignite, subbituminous coal, and other solid fuels. All values correspond to 2019 estimates..

### S. Ocean Warming

*Heat uptake* = $346 \pm 51\ TW$ *(HuID: 94108)*
*Upper ocean (0 - 700m) temperature increase since to 1960* = $0.18 - 0.20$ ℃ *(HuID: 69674, 72086)*
**Data Source(s):** Table S1 of Cheng et al. 2017. DOI: 10.1126/sciadv.1601545. NOAA National Centers for Environmental Information, 2020. DOI: 10.1029/2012GL051106.
**Notes:** Heat uptake reported is the average over time period 1992-2015 with 95% confidence intervals. Range of temperatures reported captures the 95% confidence interval of temperature increase for the period 2015-2019 with respect to the 1958-1962 mean. Temperature change is considered in the upper 700 m because sea surface temperatures have high decadal variability and are a poor indicator of ocean warming; see Roemmich et al. 2015, DOI: 10.1038/NCLIMATE2513.

### T. Power From Nuclear Fission

*Nuclear power ≈ 0.79 - 0.92 TW (HuID: 48387; 71725)*
**Data Source(s):** bp Statistical Review of World Energy, 2020. U.S. Energy Information Administration (EIA), 2022
**Notes:** Values are self-reported by countries and correspond to estimates for the 2019 calendar year.. Values are reported as 'input-equivalent' energy, meaning the energy that would have been needed to produce a given amount of power if the input were a fossil fuel, which is converted to TW here. This is calculated by

multiplying the given power by a conversion factor representing the efficiency of power production by fossil fuels. In 2017, this factor was about 40%.

### U. Nuclear Fallout

*Anthropogenic $^{239}$Pu and $^{240}$Pu from nuclear weapons ≈ 1.4 × $10^{11}$ kg / yr (HuID: 42526)*

**Data Source(s):** Table 1 in Hancock et al. 2014 doi: 10.1144/SP395.15. Fallout in activity from UNSCEAR 2000 Report on Sources and Effects of Ionizing Radiation Report to the UN General Assembly -- Volume 1.

**Notes:** The approximate mass of Plutonium isotopes $^{239}$Pu and $^{240}$Pu released into the atmosphere from the ≈ 500 above-ground nuclear weapons tests conducted between 1945 and 1980. Naturally occurring $^{239}$Pu and $^{240}$Pu are rare, meaning that nearly all contemporary labile plutonium comes from human production (Taylor 2001,doi: 10.1016/S1569-4860(01)80003-6). The total mass of radionuclides released is ≈ 3300 kg with a combined radioactive fallout of ≈ 11 PBq. These values do not represent the entire 239+240Pu globally distributed mass as it excludes non-weapons sources.

### V. Contemporary Extinction

*Animal species extinct since 1500 > 750 (HuID: 44641)*

*Plant species extinct since 1500 > 120 (HuID: 86866)*

**Data Source(s):** The IUCN Red List of Threatened Species. Version 2020-2

**Notes:** Values correspond to absolute lower-bound count of animal extinctions over the past ≈ 520 years. Of the predicted ≈ 8 million animal species, the IUCN databases catalogues only ≈ 900,000 with only ≈ 75,000 being assigned a conservation status. Representation of plants and fungi is even more sparse with only ≈ 40,000 and ≈ 285 being assigned a conservation status, respectively. The number of extinct animal species is undoubtedly higher than these reported values, as signified by an inequality symbol (>).

### W. Earth Moving

*Waste and overburden from coal mining ≈ 6.5 × $10^{13}$ kg / yr (HuID: 72899)*

*Earth moved from urbanization > 1.4 × $10^{14}$ kg / yr (HuID: 59640)*

**Data Source(s):** Supplementary table 1 of Cooper et al. 2018. DOI: doi.org/gfwfhd.

**Notes:** Coal mining waste and overburden mass is calculated given commodity-level stripping ratios (mass of overburden/waste per mass of coal resource mined) and reported values of global coal production by type. Urbanization mass is presented as a lower bound estimate of the mass of earth moved from global construction projects. This comes from a conservative estimate that the ratio of the mass of earth moved per mass of cement/concrete used in construction globally is 2:1. This value is highly context dependent and we encourage the reader to read the source material for a more thorough description of this estimation.

*Erosion rate from agriculture > (1.2 - 2.4) × $10^{13}$ kg / yr (HuID: 19415; 41496)*

**Data Source(s):** Pg. 377 of Wang and Van Oost 2019. DOI: 10.1177/0959683618816499. Pg. 21996 of Borrelli et al. 2020 DOI: 10.1073/pnas.2001403117.

**Notes:** Cumulative sediment mass loss over history of human agriculture due to accelerated erosion is estimated to be ≈ 30,000 Gt. Recent years have an estimated erosion rate ranging from 12 Pg / yr (Wang and Van Oost) to ≈ 24 Pg / yr (Borrelli et al.). Values come from computational models conditioned on time-resolved measurements of sediment deposition in catchment basins.
