## Supplementary material for "Anthroponumbers.org: A Quantitative Database Of Human Impacts on Planet Earth": Graphical Snapshot

### THE ANTHROPOCENE
#### by the numbers

Griffin Chure[1†], Rachel A. Banks[2,3,†], Avi I. Flamholz[2], Nicholas S. Sarai[4], Mason Kamb[5], Ignacio Lopez-Gomez[3,6], Yinon M. Bar-On[7], Ron Milo[7], Rob Phillips[2,5,8,*]

1. Department of Biology, Stanford University, Palo Alto, CA, USA.
California Institute of Technology, Pasadena, CA, USA, 91125:
2. Division of Biology and Biological Engineering; 3. Resnick Institute of Sustainability
4. Division of Chemistry and Chemical Engineeringt
6. Department of Environmental Science and Engineering; 8. Department of Physics
5. Chan-Zuckerberg BioHub, San Francisco, CA, 94158
7. Weizmann Institute of Science, Rehovot 7610001, Israel: Department of Plant and Environmental Sciences
†. contributed equally

#### ABSTRACT
A candidate for the greatest experiment of the last 10,000 years is the presence and action of modern human beings on planet Earth, the often complex results of which are now being felt on various fronts. While there has been a deluge of careful studies exploring each facet of these "human impacts" on Earth, they are often highly focused and necessarily technical, rarely displaying their integration with other human impacts as a whole. In this snapshot, we present a diverse (yet necessarily incomplete) array of quantities that summarize the broad reach of human action across the planet.

##### CATEGORIES
- LAND
- ENERGY
- WATER
- FLORA & FAUNA
- ATMOSPHERIC & BIOGEOCHEMICAL CYCLES

##### REFERENCE UNITS
human population ≈ $8\times10^9$

**area**: soccer field ≈ 7000 m²; tennis court ≈ 300 m²; room ≈ 10 m²

**volume**: shipping container ≈ 30 m³; bathtub ≈ 0.1 m³

**power**: incandescent lightbulb ≈ 100 W

**mass**: pick-up truck ≈ 2000 kg; refrigerator ≈ 100 kg; cinder block ≈ 10 kg

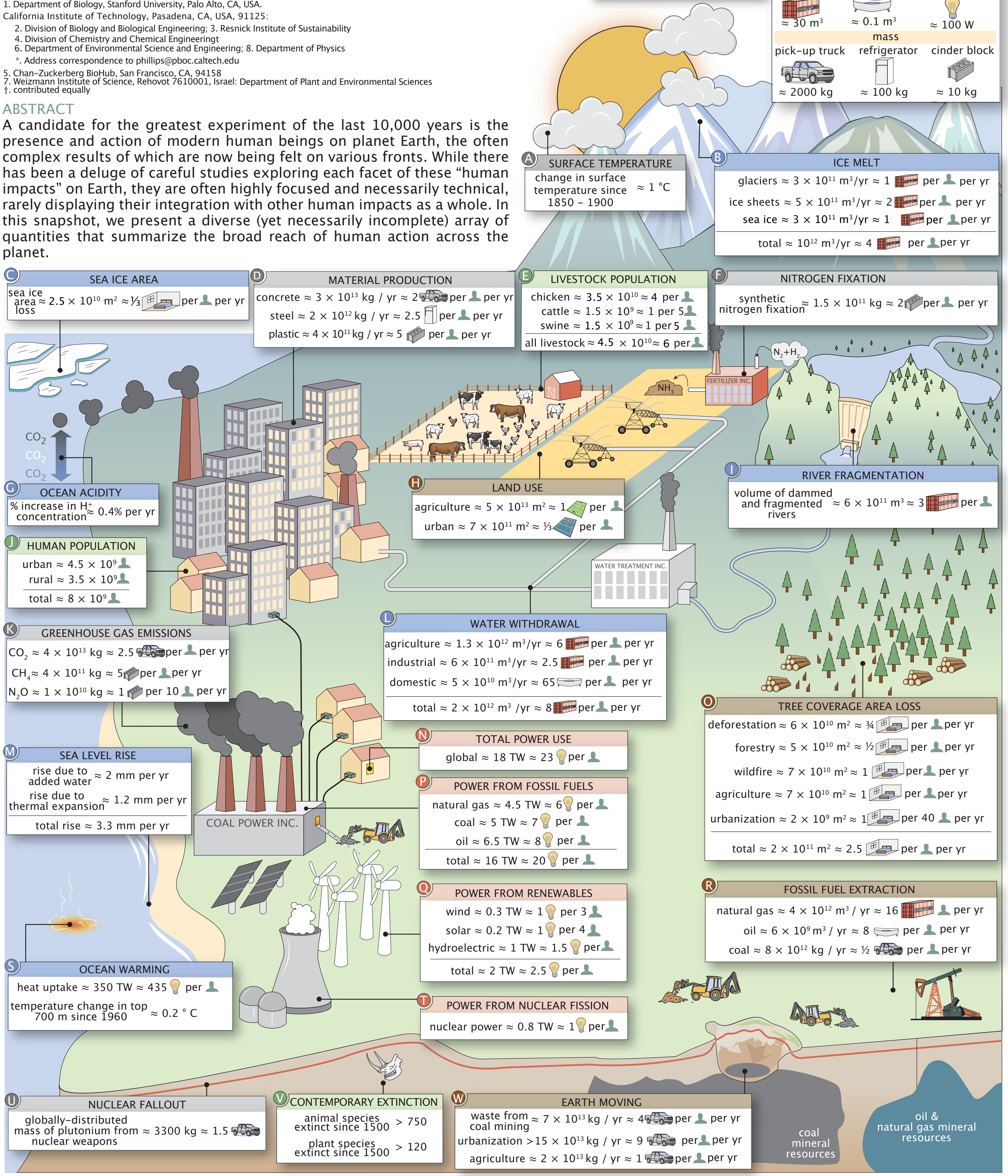


##### A SURFACE TEMPERATURE
change in surface temperature since 1850 – 1900 ≈ 1 °C

##### B ICE MELT
glaciers ≈ $3\times10^{11}$ m³/yr ≈ 1 per per yr
ice sheets ≈ $5\times10^{11}$ m³/yr ≈ 2 per per yr
sea ice ≈ $3\times10^{11}$ m³/yr ≈ 1 per per yr
total ≈ $10^{12}$ m³/yr ≈ 4 per per yr

##### C SEA ICE AREA
sea ice area loss ≈ $2.5\times10^{10}$ m² ≈ ⅓ per per yr

##### D MATERIAL PRODUCTION
concrete ≈ $3\times10^{13}$ kg / yr ≈ 2 per per yr
steel ≈ $2\times10^{12}$ kg / yr ≈ 2.5 per per yr
plastic ≈ $4\times10^{11}$ kg / yr ≈ 5 per per yr

##### E LIVESTOCK POPULATION
chicken ≈ $3.5\times10^{10}$ ≈ 4 per
cattle ≈ $1.5\times10^9$ ≈ 1 per 5
swine ≈ $1.5\times10^9$ ≈ 1 per 5
all livestock ≈ $4.5\times10^{10}$ ≈ 6 per

##### F NITROGEN FIXATION
synthetic nitrogen fixation ≈ $1.5\times10^{11}$ kg ≈ 2 per per yr

##### G OCEAN ACIDITY
% increase in $H^+$ concentration ≈ 0.4% per yr

##### H LAND USE
agriculture ≈ $5\times10^{13}$ m² ≈ 1 per
urban ≈ $7\times10^{11}$ m² ≈ ⅓ per

##### I RIVER FRAGMENTATION
volume of dammed and fragmented rivers ≈ $6\times10^{11}$ m³ ≈ 3 per

##### J HUMAN POPULATION
urban ≈ $4.5\times10^9$
rural ≈ $3.5\times10^9$
total ≈ $8\times10^9$

##### K GREENHOUSE GAS EMISSIONS
$CO_2$ ≈ $4\times10^{13}$ kg ≈ 2.5 per per yr
$CH_4$ ≈ $4\times10^{11}$ kg ≈ 5 per per yr
$N_2O$ ≈ $1\times10^{10}$ kg ≈ 1 per 10 per yr

##### L WATER WITHDRAWAL
agriculture ≈ $1.3\times10^{12}$ m³/yr ≈ 6 per per yr
industrial ≈ $6\times10^{11}$ m³/yr ≈ 2.5 per per yr
domestic ≈ $5\times10^{10}$ m³/yr ≈ 65 per per yr
total ≈ $2\times10^{12}$ m³ /yr ≈ 8 per per yr

##### M SEA LEVEL RISE
rise due to added water ≈ 2 mm per yr
rise due to thermal expansion ≈ 1.2 mm per yr
total rise ≈ 3.3 mm per yr

##### N TOTAL POWER USE
global ≈ 18 TW ≈ 23 per

##### O TREE COVERAGE AREA LOSS
deforestation ≈ $6\times10^{10}$ m² ≈ ¾ per per yr
forestry ≈ $5\times10^{10}$ m² ≈ ½ per per yr
wildfire ≈ $7\times10^{10}$ m² ≈ 1 per per yr
agriculture ≈ $7\times10^{10}$ m² ≈ 1 per per yr
urbanization ≈ $2\times10^9$ m² ≈ 1 per 40 per yr
total ≈ $2\times10^{11}$ m² ≈ 2.5 per per yr

##### P POWER FROM FOSSIL FUELS
natural gas ≈ 4.5 TW ≈ 6 per
coal ≈ 5 TW ≈ 7 per
oil ≈ 6.5 TW ≈ 8 per
total ≈ 16 TW ≈ 20 per

##### Q POWER FROM RENEWABLES
wind ≈ 0.3 TW ≈ 1 per 3
solar ≈ 0.2 TW ≈ 1 per 4
hydroelectric ≈ 1 TW ≈ 1.5 per
total ≈ 2 TW ≈ 2.5 per

##### R FOSSIL FUEL EXTRACTION
natural gas ≈ $4\times10^{12}$ m³ / yr ≈ 16 per yr
oil ≈ $6\times10^9$ m³ / yr ≈ 8 per per yr
coal ≈ $8\times10^{12}$ kg / yr ≈ ½ per per yr

##### S OCEAN WARMING
heat uptake ≈ 350 TW ≈ 435 per
temperature change in top 700 m since 1960 ≈ 0.2 ° C

##### T POWER FROM NUCLEAR FISSION
nuclear power ≈ 0.8 TW ≈ 1 per

##### U NUCLEAR FALLOUT
globally-distributed mass of plutonium from nuclear weapons ≈ 3300 kg ≈ 1.5

##### V CONTEMPORARY EXTINCTION
animal species extinct since 1500 > 750
plant species extinct since 1500 > 120

##### W EARTH MOVING
waste from coal mining ≈ $7\times10^{13}$ kg / yr ≈ 4 per per yr
urbanization > $15\times10^{13}$ kg / yr ≈ 9 per per yr
agriculture ≈ $2\times10^{13}$ kg / yr ≈ 1 per per yr

### The Anthropocene by the Numbers — Impacts By Region

#### A THE GEOGRAPHY OF HUMAN IMPACTS

Page 1 represents the impact humans have on the Earth at a global scale. While these numbers are handy, it is important to acknowledge that they vary from country-to-country and continent-to-continent. Furthermore, the consequences of these anthropogenic impacts are also unequally distributed, meaning some regions experience effects disproportionate to their contribution. Here, we give a sense of the geographic distribution of several values presented on page 1, broken down by continental region as shown below.

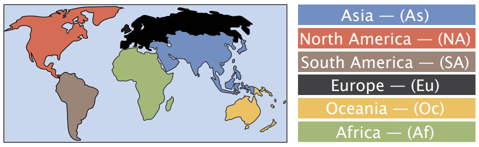


#### B THE HUMAN POPULATION

There are ≈ 8 billion humans on the planet, with approximately 50% living in 'urban' environments. The majority of the worlds population (as well as the majority of both urban and rural dwellers) live in Asia.

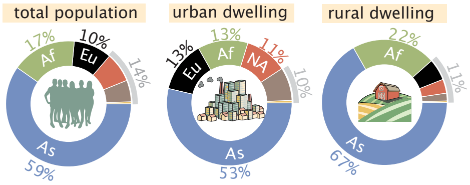


Sources: Food and Agricultural Organization of the United Nations – World Population
Notes: Urban/rural designation has no set definition and follows the conventions set by each reporting country.

#### C LAND USE

Though humans are nearly evenly split between urban and rural environments, agricultural land is the far more common use of land area. Together, Asia and Africa contain more than half of global agricultural land. Asia alone accomodates more than half of the global urban land area.

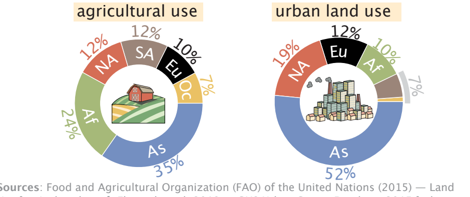


Sources: Food and Agricultural Organization (FAO) of the United Nations (2015) — Land Use [agricultural area]. Florczyk et al. 2019 — GHS Urban Centre Database 2015 [urban land area] Notes: Urban is defined as any inhabited area with ≥ 2500 residents, as defined by the USDA.

#### D THE LIVESTOCK POPULATION

The global population of terrestrial livestock is around 30 billion individuals, most of which are chickens. Asia houses most of the global livestock population, though South America and Europe harbor more animals on a per-capita basis.

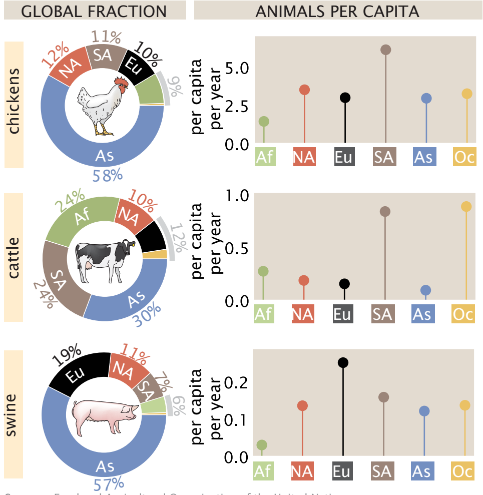


Sources: Food and Agricultural Organization of the United Nations

#### E WATER WITHDRAWAL

While Asia withdraws the most water for agricultural and municipal needs, North America withdraws the plurality of water for industrial purposes. North America also withdraws more water per capita than any other region.

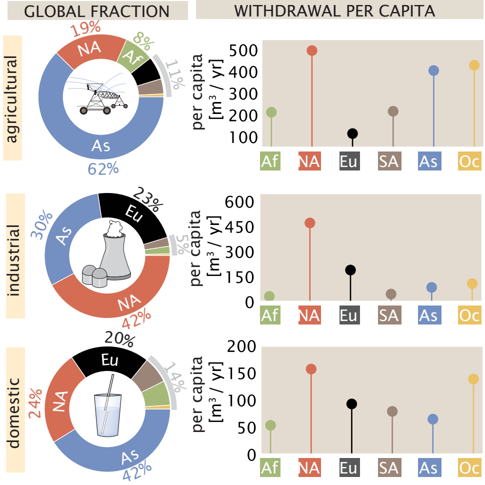


Source: AQUASTAT Main Database, Food and Agriculture Organization of the United Nations.
Notes: Values are reported directly from member countries and represent average of 2013–2017 period. Per capita values are computed given population of reporting countries.

#### F TREE COVERAGE AREA LOSS

Most drivers of tree coverage area loss are comparable in their effect at a global scale. However, there are drastic regional differences in the relative magnitudes.

REGION DEFINITION

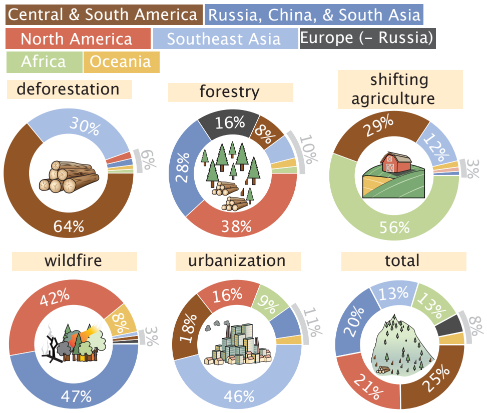


Source: Curtis et al. 2018 doi: 10.1126/science.aau3445.
Notes: Regions are as reported in Curtis et al. 2018. "Deforestation" here denotes permanent removal of tree cover for commodity production. "Shifting agriculture" here denotes forest/shrub land converted to agriculture and later abandoned. All values correspond to breakdown of cumulative tree cover area loss from 2001 – 2015.

#### G NITROGENOUS FERTILIZER USE & PRODUCTION

Modern agriculture requires nitrogen in amounts beyond what is produced naturally. Asia synthesizes and consumes a large majority of fixed nitrogen. However, Europe and North America dominate per capita synthesis whereas Oceania consumes more fertilizer per capita than any other region.

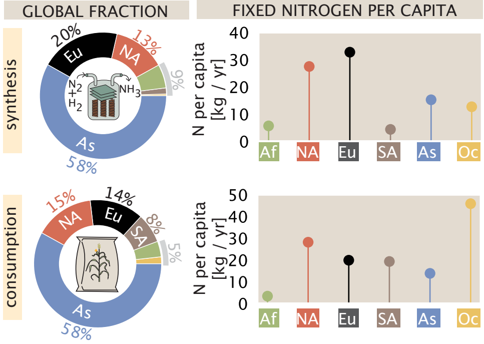


Source: Food and Agricultural Organization (FAO) of the United Nations.
Notes: Values account for reactive nitrogen production/consumption in context of fertilizer only and does not account for plastics, explosives, or other uses.

#### H GREENHOUSE GAS EMISSIONS

$CO_2$ and $CH_4$ are two potent greenhouse gases which are routinely emitted by anthropogenic processes such as burning fuel and rearing livestock. While Asia emits roughly half of all $CO_2$ and $CH_4$, North America and Oceania produce the most on a per capita basis, respectively.

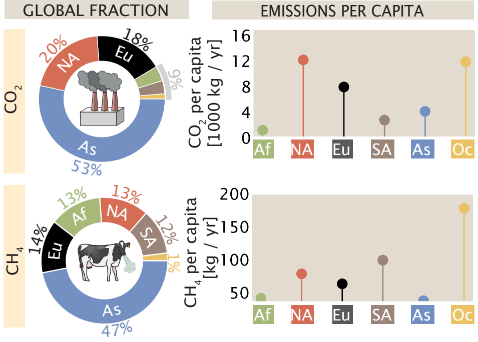


Sources: $CO_2$ data collated by: Friedlingstein, P. et al. (2019). doi: 10.5194/essd-11-1783-2019. See Panel K on Pg. 4 for complete list of sources. $CH_4$ data from Saunois et al, 2020 doi: 10.5194/essd-12-1561-2020 Notes: Values report decadal averages in kg $CO_2$ or $CH_4$ per year over time period 2008–2017.

#### I MATERIAL PRODUCTION

Humans excavate an enormous amount of material from the Earth's crust and transform it to build our structures. Two of these materials, concrete and steel, are produced primarily in Asia on both a global and per capita basis. Asia's per capita production of steel is only outpaced by Europe.

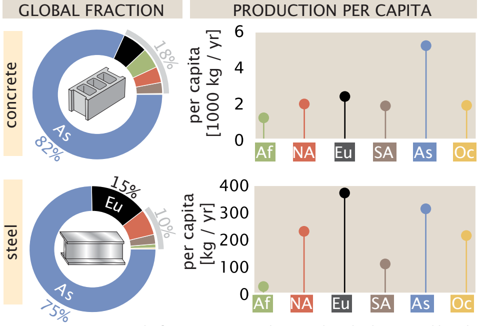


Sources: USGS Statistics and Information 2020, Steel Statistical Yearbook 2019 World Steel Association. Food and Agricultural Organization (FAO) of the United Nations — Annual Population. Notes: Reported values for cement and steel production corresponds to 2017 and 2018 values, respectively. Mass of concrete was calculated using a rule-of-thumb that 1 kg of cement yields 7 kg of concrete (Monteiro et al. 2017. doi: 0.138/nmat4930).

#### J POWER GENERATION AND CONSUMPTION

From heating water, to powering lights, to moving our vehicles, nearly every facet of modern human life requires the consumption of power, culminating in nearly 20 TW of power use in recent years. Asia consumes over half of the power derived from combustion of fossil fuels, with Europe and North America each consuming around 20% of the global total. Asia also produces the plurality of power from renewable technologies, such as hydroelectric, wind, and solar, however, North America, South America, and Europe each produce more on a per capita basis. Nuclear energy, however, is primarly produced in Europe, with North America and Asia coming in second and third place, respectively. On a per-capita basis, North America consumes or produces more energy than all other regions considered here, yielding a total power consumption of nearly 10,000 W per person.

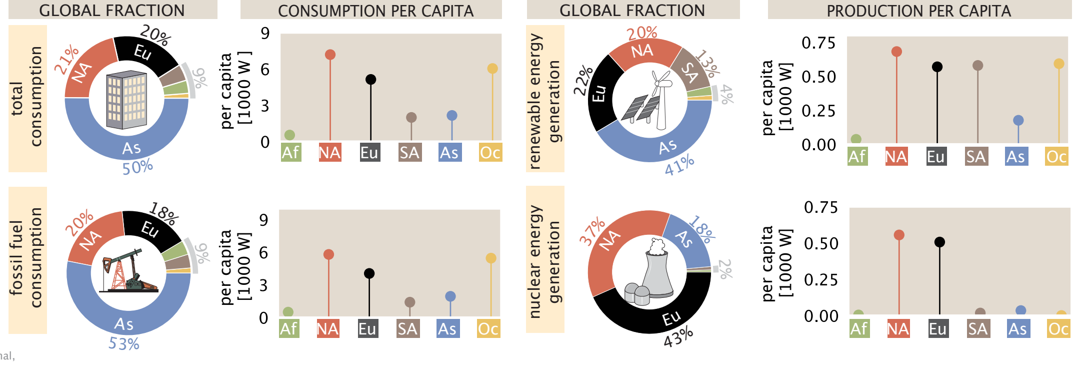


Source: Energy Information Administration of the United States (2017)
Notes: "Renewables" includes hydroelectric, biofuels, biomass (wood), geothermal, wind, and solar. "Fossil fuels" includes coal, oil, and natural gas.

### The Anthropocene by the Numbers — Dynamics of Global Magnitudes

#### A THE HUMAN POPULATION

The human population has more than doubled in the past 60 years. During this time, the fraction of the population living in urban areas has increased so that today the population is about evenly split between urban and rural.

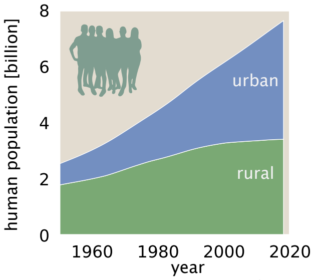


**Sources:** Food and Agricultural Organization of the United Nations – World Population
**Notes:** Urban/rural designation has no set definition and follows the conventions set by each reporting country.

#### B WATER WITHDRAWAL

Total water withdrawal has increased with the human population, dominated by increasing agricultural use. Per capita, the amount of water withdrawn for agriculture and domestic use has remained constant since the 1980s.

##### ANNUAL WATER WITHDRAWAL

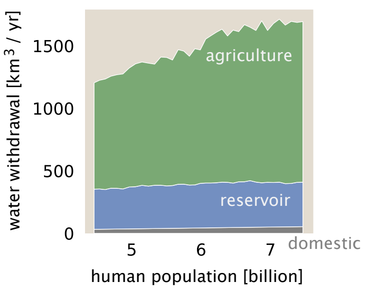


##### WITHDRAWAL PER CAPITA

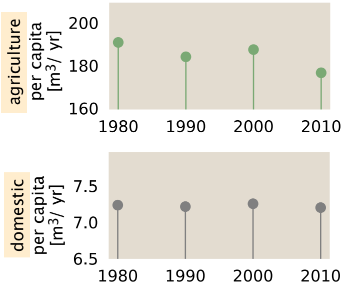


**Source:** AQUASTAT Main Database, Food and Agriculture Organization of the United Nations.
**Notes:** Values are reported directly from member countries and represent average of 2013-2017 period. Per capita values are computed given population of reporting countries.

#### C THE LIVESTOCK POPULATION

The standing population of livestock has been increasing, with chicken making up a large fraction of the total livestock population. The number of chicken raised per capita has increased since the 1960s, while cattle per capita have decreased.

##### STANDING POPULATION

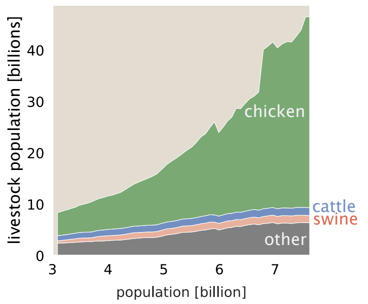


##### LIVESTOCK PER CAPITA

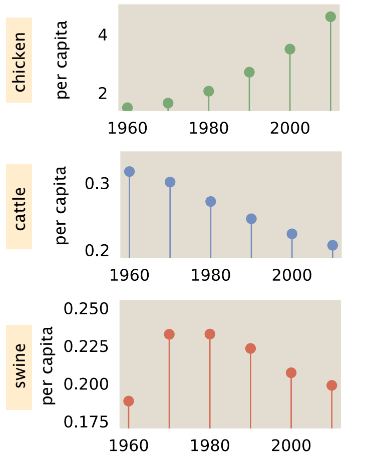


**Sources:** Food and Agricultural Organization of the United Nations

#### D MATERIAL PRODUCTION

Total mass of human-made materials hasbeen accumulating over time, dominated byconstruction materials. Per capita, the massof bricks & asphalt, aggregates, andconcrete has increased since the 1950s.

##### TOTAL ANTHROPOMASS

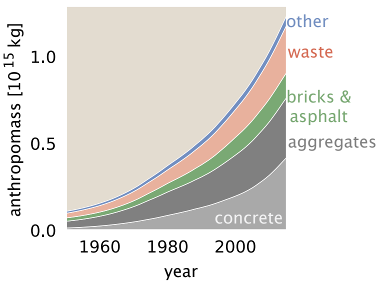


##### MASS PER CAPITA

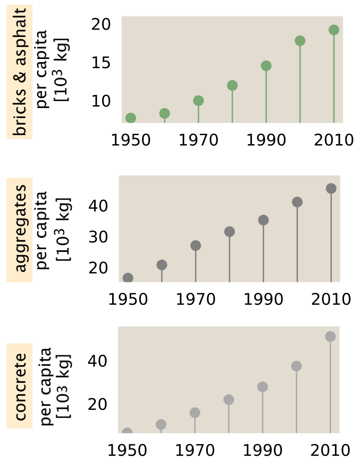


**Sources:** Krausmann et al. 2017 doi: 10.1073/pnas.1613773114
**Notes:** Material production is estimated from a material flows model.

#### E AQUATIC FOODS PRODUCTION

Aquatic (blue) foods production has been increasing with the human population. Interestingly, the mass produced from wild capture has remained constant per capita since the 1980s while the mass produced by aquaculture has increased per capita during the same period, driving the increase in overall production.

##### ANNUAL BLUE FOOD PRODUCTION

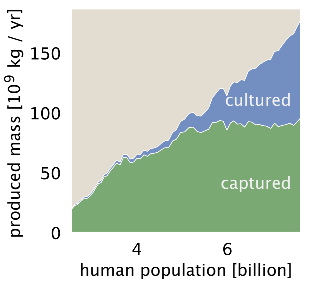


##### PRODUCTION PER CAPITA

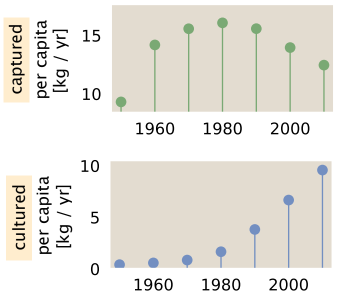


**Sources:** Food and Agricultural Organization of the United Nations

#### F POWER CONSUMPTION

Power consumption has increased with population, as well as technological and societal changes, which have driven an increase in power per capita across all generation types. The source of our power has also changed over time. Over the last 60 years, nuclear power has become comparable to hydroelectricity, with most of the growth occurring between 1970 and 1990. Renewable power generation is currently experiencing a similar growth pattern.

##### ANNUAL POWER CONSUMPTION

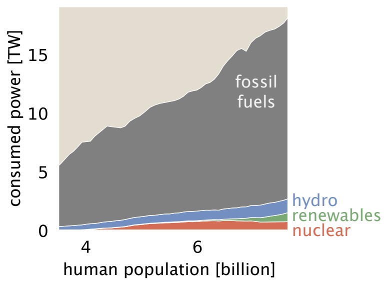


##### CONSUMPTION PER CAPITA

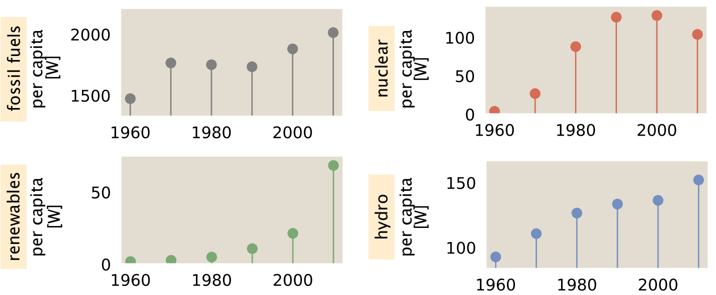


**Source:** Energy Information Administration of the United States (2017)
**Notes:** "Renewables" includes biofuels, biomass (wood), geothermal, wind, and solar. "Fossil fuels" includes coal, oil, and natural gas.

#### G $CO_2$ EMISSIONS

Annual anthropogenic $CO_2$ emissions have been increasing with the population, driven by an increase in fossil fuel combustion. The amount of $CO_2$ emissions from fossil fuels has increased slightly per capita, while the per capita emissions from land use change have decreased.

##### ANNUAL $CO_2$ EMISSIONS

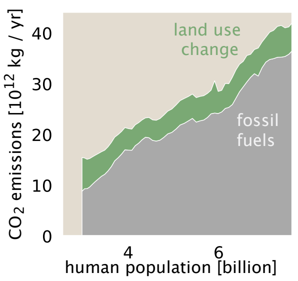


##### EMISSIONS PER CAPITA

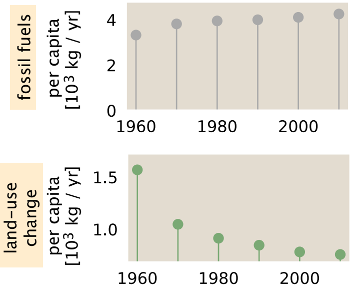


**Sources:** Data collated by: Friedlingstein, P. et al. (2019). doi: 10.5194/essd-11-1783-2019.
See Panel K on Pg. 4 for complete list of sources

#### H $CH_4$ EMISSIONS

While total anthropogenic methane ($CH_4$) emissions have been increasing with the human population, per capita emissions have been decreasing each decade since the 1970s. This per capita emission reduction reflects a shift in global diets away from methane-intenesive beef products, as well as better waste management policies in developed countries.

##### ANNUAL $CH_4$ EMISSIONS

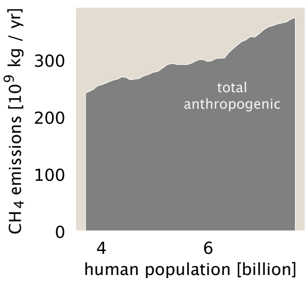


##### EMISSIONS PER CAPITA

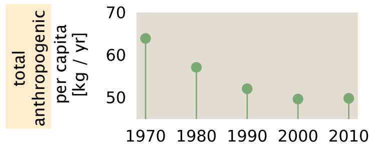


**Sources:** Saunois et al, 2020 doi: 10.5194/essd-12-1561-2020

### The Anthropocene by the Numbers — Supporting Information

About: Here, we present citations and notes corresponding to each quantity assessed here. Each value presented on page 1 is assigned a Human Impacts Database identifier (HuID), accessible via anthroponumbers.org. When possible, primary data sources have been collated and stored as files in comma-separated-value (csv) format on the GitHub repository associated with this snapshot, accessible via DOI: 10.5281/zenodo.4453277 and https://github.com/rpgroup-pboc/human_impacts

#### A SURFACE TEMPERATURE

Surface temperature change relative to 1850-1900 average ≈ 1 - 1.4° C — HuID: 79598, 76539, 12147

Data Source(s): HadCRUT.4.6 (Morice et al., 2012, DOI: 10.1029/2011JD017187), GISTEMP v4 (GISTEMP Team, 2020: GISS Surface Temperature Analysis (GISTEMP), version 4. NASA Goddard Institute for Space Studies. Dataset accessed 2020-12-17 at https://data.giss.nasa.gov/gistemp/ & Lenssen et al., 2019, DOI: 10.1029/2018JD029522) and NOAAGlobalTemp v5 (Zhang et al, 2019, DOI: 10.1029/2019EO128229) datasets. Notes: The global mean surface temperature captures near-surface air temperature over the planet's land and ocean surface. The value reported represents the spread of the three estimates and their 95% confidence intervals. Temperature changes from all three datasets are expressed relative to the 1850-1900 average temperature from the HadCRUT.4.6 dataset. Since data for the period 1850-1880 are missing in GISTEMP v4 and NOAAGlobalTemp v5, data are centered by setting the 1880-1900 mean of all datasets to the HadCRUT.4.6 mean over the same period.

#### E LIVESTOCK POPULATION

| | |
|---|---|
| chicken standing population ≈ 3.5 × $10^{10}$ | HuID: 94934 |
| cattle standing population ≈ 1.5 × $10^9$ | HuID: 92006 |
| swine standing population ≈ 1.5 × $10^9$ | HuID: 21368 |
| all livestock standing population ≈ 4.6× $10^{10}$ | HuID: 43599 |

Data Source(s): Food and Agriculture Organization (FAO) of the United Nations Statistical Database (2020) — Live Animals. Notes: Counts correspond to the estimated standing populations in 2019. Values are reported directly by countries. The FAO uses non-governmental statistical sources to address uncertainty and missing (non-reported) data. Reported values are therefore approximations.

#### F SYNTHETIC NITROGEN FIXATION

annual mass of synthetically fixed nitrogen ≈ (1.4 - 1.5)× $10^{11}$ kg N / yr — HuID: 60580, 61614

Data Source(s):United States Geological Survey (USGS) National Minerals Information Center, Commodity Statistics and Information, Nitrogen Statistics and Information International Fertilizer Association (IFA) Statistical Database (2021) Smith et al. 2020DOI:10.1039 /c9ee02873k Notes: Ammonia ($NH_3$) produced globally is compiled by the USGS and IFA from major factories that report output. The USGS estimates the approximate mass of nitrogen in ammonia produced in 2019 as 1.42 ×$10^{11}$ kg N and the International Fertilizer Association reports a production value of 1.50 ×$10^{11}$ kg N in 2019. Nearly all of this mass is produced by the Haber-Bosch process (>96%, Smith et al. 2020). In the United States, most of this mass is used for fertilizer, with the remainder being used to synthesize nitrogen-containing chemicals including explosives, plastics, and (≈ 88%, USGS Mineral Commodity Summaries 2020 - Nitrogen

#### G OCEAN ACIDITY

| | |
|---|---|
| surface ocean [H+] ≈ 0.2 parts per billion | HuID: 90472 |
| annual change in [H+] = 0.36 ± 0.03 % | HuID: 19394 |

Data Source(s): Figures 1-2 of European Environment Agency report CLIM 043 (2020). Original data source of the report is "Global Mean Sea Water pH" from Copernicus Marine Environment Monitoring Service. Notes: Reported value is calculated from the global average annual change in pH over years 1985-2018. The average oceanic pH was ≈ 8.057 in 2018 and decreases annually by ≈ 0.002 units, giving a change in [H+] of roughly $10^{-8.056}$ - $10^{-8.057}$ ≈ 4×$10^{-11}$ mol/L or about 0.4% of the global average. [H+] is calculated as $10^{-pH}$ ≈ $10^{-8}$ mol/L or 0.2 parts per billion (ppb) which is calculated by noting that [$H_2O$] ≈ 55 mol / L. Uncertainty for annual change is the standard error of the mean.

#### L WATER WITHDRAWAL

| | |
|---|---|
| agricultural withdrawal = 1.3 × $10^{12}$ m³ / yr | HuID: 84545, 43593, 95345 |
| industrial withdrawal = 5.9 × $10^{11}$ m³ / yr | HuID: 27142 |
| domestic withdrawal = 5.4 × $10^{10}$ m³ / yr | HuID: 69424 |
| total withdrawal = (1.7 – 2.2) × $10^{12}$ m³ / yr | HuID: 27342, 68004 |

#### O TREE COVERAGE AREA LOSS

| | |
|---|---|
| commodity-driven deforestation = (5.7 ± 1.1) × $10^{10}$ m² / yr | HuID: 96098 |
| forestry = (5.4 ± 0.8) × $10^{10}$ m² / yr | HuID: 38352 |
| urbanization = (2 ± 1) × $10^{9}$ m² / yr | HuID: 19429 |
| shifting agriculture = (7.5 ± 0.9) × $10^{10}$ m² / yr | HuID: 24388 |
| wildfire = (7.2 ± 1.3) × $10^{10}$ m² / yr | HuID: 92221 |
| total loss ≈ 2 × $10^{11}$ m² / yr | HuID: 78576 |

Data Source(s): Table 1 of Curtis et al. 2018 DOI:10.1126/science.aau3445. Hansen et al. 2013 DOI:10.1126/science.1244693. Global Forest Watch, 2020. Reported values in source correspond to total loss from 2001 – 2015. Values given are averages over this 15 year window. Notes: Commodity-driven deforestation is "long-term, permanent, conversion of forest and shrubland to a non-forest land use such as agriculture, mining, or energy infrastructure." Forestry is defined as large-scale operations occurring within managed forests and tree plantations with evidence of forest regrowth in subsequent years. Urbanization converts forest and shrubland for the expansion and intensification of existing urban centers. Disruption due to "shifting agriculture" is defined as "small- to medium-scale forest and shrubland conversion for agriculture that is later abandoned and followed by subsequent forest regrowth". Disruption due to wildfire is "large-scale forest loss resulting from the burning of forest vegetation with no visible human conversion or agricultural activity afterward". Uncertainty corresponds to the 95% confidence interval. Uncertainty is approximate for "urbanization" as the source reports an ambiguous error of "± <1%".

#### P POWER FROM FOSSIL FUELS

| | |
|---|---|
| natural gas = 4.5 – 4.9 TW | HuID: 49947, 86175 |
| oil = 6.1 – 6.6 TW | HuID: 4121, 39756 |
| coal = 5.0 – 5.6 TW | HuID: 10400, 60490 |
| total = 16 – 17 TW | HuID: 29470, 29109 |

Data Source(s): bp Statistical Review of World Energy, 2022. U.S. Energy Information Administration, 2020. Notes: Values are self-reported by countries. values from bp Statistical Review and EIA correspond to 2019. Reported TW are computed from primary energy (e.g. kg coal) units assuming uniform use throughout the year. Oil volume includes crude oil, shale oil, oil sands, condensates, and natural gas liquids separate from specific natural gas mining. Natural gas value excludes gas flared or recycled and includes natural gas produced for gas-to-liquids transformation. Coal value includes 2019 value exclusively for solid commercial fuels such as bituminous coal and anthracite, lignite and subbituminous coal, and other solid fuels. This includes coal used directly in power production as well as coal used in coal-to-liquids and coal-to-gas transformations.

#### Q POWER FROM RENEWABLE RESOURCES

| | |
|---|---|
| wind ≈ 0.36– 0.43 TW | HuID: 30581, 85919 |
| solar ≈ 0.18 – 0.21 TW | HuID: 99885, 58303 |
| hydroelectric = 1.2 TW | HuID: 15765, 50558 |
| total renewable power ≈ 1.9 – 2.1 TW | HuID: 75741, 20246 |

Data Source(s): bp Statistical Review of World Energy, 2022. U.S. Energy Information Administration, 2020. Notes: Reported values correspond to estimates for the 2019 calendar year. Renewable resources are defined as wind, geothermal, solar, biomass and waste. Hydroelectric, while presented here, is not defined as a renewable in the bp dataset. All values are reported as input-equivalent energy, meaning the input energy that would have been required if the power was produced by fossil fuels. bp reports that fossil fuel efficiency used to make this conversion was about 40% in 2017.

#### R FOSSIL FUEL EXTRACTION

| | |
|---|---|
| volume of natural gas = (3.9 – 4.0) × $10^{12}$ m³ / yr | HuID: 11468, 20532 |
| volume of oil = (5.5 ± 5.8) × $10^{9}$ m³ / yr | HuID: 66789, 97719 |
| mass of coal = (7.8 – 8.1) × $10^{12}$ kg / yr | HuID: 78435, 48928 |

Data Source(s): bp Statistical Review of World Energy, 2022. U.S. Energy Information Administration, 2020. Notes: Oil volume includes crude oil, shale oil, oil sands, condensates, and natural gas liquids separate from specific natural gas mining. Natural gas value excludes gas flared or recycled and includes natural gas produced for gas-to-liquids transformation. Coal value includes solid commercial fuels such as bituminous coal, anthracite, lignite, subbituminous coal, and other solid fuels. All values correspond to 2019 estimates.

We are incredibly grateful for the generosity of a wide array of experts for their advice, suggestions, and criticism of this work. Specifically, we thank Suzy Beeler, Lars Bildsten, Justin Bois, Chris Bowler, Matthew Burgess, Ken Caldeira, Jörn Callies, Sean B. Carroll, Ibrahim Cissé, Joel Cohen, Michelle Dan, Bethany Ehlmann, Gidon Eshel, Paul Falkowski, Daniel Fisher, Thomas Frederikse, Neil Fromer, Eric Galbraith, Lea Goentoro, Evan Groover, John Grotzinger, Soichi Hirokawa, Greg Huber, Christina Hueschen, Bob Jaffe, Elizabeth Kolbert, Thomas Lecuit, Raphael Magarik, Jeff Marlow, Brad Marston, Jitu Mayor, Elliot Meyerowitz, Lisa Miller, Dianne Newman, Luke Oltrogge, Nigel Orme, Victoria Orphan, Marco Pasti, Pietro Perona, Noam Prywes, Stephen Quake, Hamza Raniwala, Manuel Razo-Mejia, Thomas Rosenbaum, Benjamin Rubin, Alex Rubinsteyn, Shyam Saladi, Tapio Schneider, Murali Sharma, Alon Shepon, Arthur Smith, Matthieu Talpe, Wati Taylor, Julie Theriot, Tadashi Tokieda, Cat Triandifillou, Sabah Ul-Hasan, Tine Valencic, and Ned Wingreen. We also thank Yue Qin for sharing data related to global water consumption. Many of the topics in this work began during the Applied Physics 150C course taught at Caltech during the early days of the COVID-19 pandemic. This work was supported by the Resnick Sustainability Institute at Caltech and the Schwartz-Reisman Collaborative Science Program at the Weizmann Institute of Science.
